# Loss of the Er81 Transcription Factor in Cholinergic Cells Alters Striatal Activity and Habit Formation

**DOI:** 10.1101/2020.01.14.905497

**Authors:** Yadollah Ranjbar-Slamloo, Noorya Yasmin Ahmed, Alice Shaam Al Abed, Lingxiao Gao, Yovina Sontani, Alexandre R’Com-H Cheo Gauthier, Ehsan Arabzadeh, Nathalie Dehorter

## Abstract

The finely-tuned activity of cholinergic interneurons (CINs) in the striatum is key for motor control, learning, and habit formation. Yet, the molecular mechanisms that determine their unique functional properties remain poorly explored. Using a combination of genetic and biochemical assays, *in vitro* and *in vivo* physiological characterisation, we report that selective ablation of the Er81 transcription factor leads to prominent changes in CIN molecular, morphological and electrophysiological features. In particular, the lack of Er81 amplifies intrinsic delayed-rectifier and hyperpolarization-activated currents, which subsequently alters the tonic and phasic activity of CINs. We further demonstrate that these alterations enhance their pause and time-locked responses to sensorimotor inputs in awake mice. Finally, this study reveals an Er81-dependent developmental mechanism in CINs essential for habit formation in adult mice.

**Highlights:** - The Er81 transcription factor is expressed in striatal cholinergic interneurons (CINs)

- Conditional deletion of Er81 alters key molecular, morphological and electrophysiological properties of CINs in adult mice

- Deletion of Er81 reduces the intrinsic excitability of CINs by upregulating delayed rectifier and hyperpolarization-activated currents

- Deletion of Er81 alters *in vivo* striatal activity and habit formation

## INTRODUCTION

Cholinergic interneurons (CINs) constitute only 1–2% of all striatal neurons but are the main source of acetylcholine in the striatum and play a crucial role in regulating motor learning and behavioural flexibility (Aoki et al., 2015) (Aoki et al., 2018) (Martos et al., 2017). Unique morphological and electrophysiological features (Lim et al., 2014) underpin CIN function in controlling the striatal output neurons (Mamaligas and Ford, 2016) (Gritton et al., 2019) (Morris et al., 2004). CINs fire tonically and display phasic responses to stimuli, which consist of pauses preceded by a transient rise and followed by a ‘‘rebound’’of CIN activity (Apicella, 2017). The pause in CIN firing is fundamental for striatal processing of information and behaviour (Zucca et al., 2018). The tonic and phasic activity of CINs are governed by both synaptic inputs and the intrinsic inward (Zhao et al., 2016) and delayed rectifier currents (Wilson and Goldberg, 2006). In particular, pause expression is mediated by the slow Kv7-mediated potassium current I_Kr_ in response to excitatory inputs and is regulated by dopamine (Zhang et al., 2018). Yet, the role of intrinsic molecular factors in regulating CIN pauses remains unexplored.

Recent findings suggest that developmental differentiation factors induce some degree of functional diversity amongst CINs, thus enabling them to acquire unique properties (Magno et al., 2017) (Lozovaya et al., 2018) (Ahmed et al., 2019). However, the functional relevance of this heterogeneity remains unclear. The ETS transcription factor Er81 plays a crucial role in cell maturation (Abe et al., 2011) (Willardsen et al., 2014) (Ding et al., 2016), identity (Doitsidou et al., 2013) (Cave et al., 2010) (Flames and Hobert, 2009), excitability (Dehorter et al., 2015), and the establishment of synaptic connections (Arber et al., 2000) (Hippenmeyer et al., 2005). It is expressed in the striatum (Nobrega-Pereira et al., 2008) (Mi et al., 2018), but its specific function is unknown. We hypothesised that Er81 plays a fundamental role in determining key features of developing CINs and in controlling striatal CIN function. Using cell-type specific gene ablation, molecular assays, together with *in vitro* and *in vivo* physiological characterisation, we reveal that the Er81 transcription factor is necessary to set major functional properties of CINs. We unravel its role as a key contributing factor for pause expression, regulation of striatal activity and habitual behaviour.

## RESULTS

### Er81 is expressed in the striatal cholinergic interneurons

We first analysed the expression of Er81 from birth to adulthood and found that both mRNA (**Figure 1A**) and protein levels (**Figure 1B**) significantly drop from postnatal day 6 (P6) to P30 in the total striatum. In the CINs specifically, we observed similar decrease of Er81 protein levels from P6 to P30 (**Figure 1C**). To determine the proportion of CINs expressing Er81 at any stage during development, we analysed β-galactosidase staining, which perdures long after being synthesised (Dehorter et al., 2015). We found that most of CINs (62 ± 7 %) in the Er81*^nlsLacZ/+^* mice express β-galactosidase (and therefore Er81) in the striatum (**Figure S1A)**, unlike cholinergic cells in the basal forebrain (6 ± 2%; **Figure S1B**). As Er81 can have an important role in cell function (Dehorter et al., 2015), we analysed the consequences of removing *Er81* from CINs, using conditional knockout mice (cKO: *ChAT-Cre; Er81^f/f^*; comparing with control mice: *ChAT-Cre; Er81^+/+^*). We did not observe any change in cholinergic cell density at P2 and P30 within the striatum between control and *Er81* cKO conditions (**Figure 1D-F**), suggesting that the specific deletion of *Er81* does not affect CIN neurogenesis and migration.

**Figure 1:**
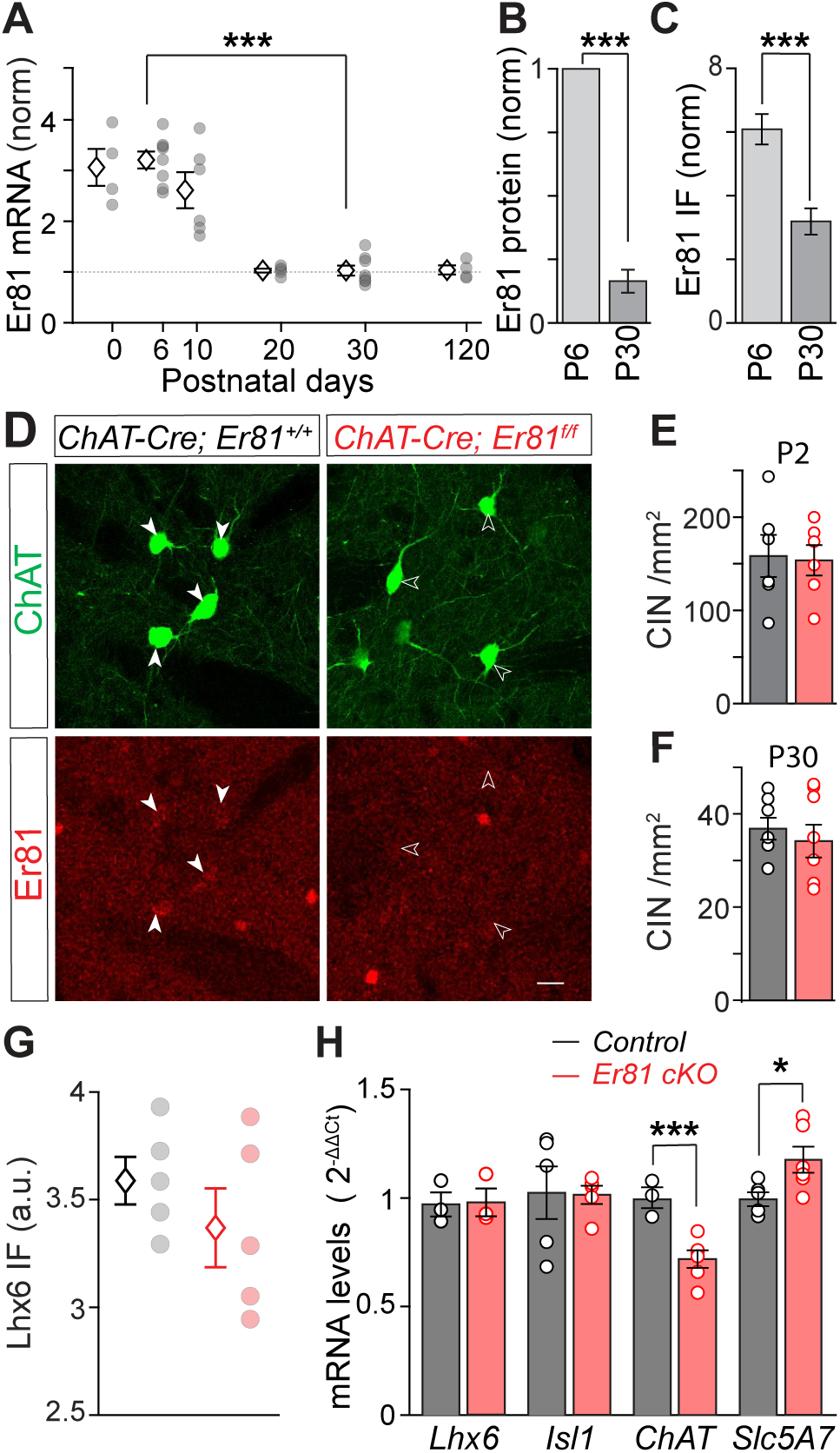
Er81 is expressed in the developing striatum and in cholinergic interneurons. **A.** *Er81* mRNA levels from P0 to P120 dorsal striatum (n = 4-8 mice). Data normalised to P30. **B.** Er81 protein levels from P6 (n = 4 mice) and P30 striatum (n = 6 mice, normalised to P6, one sample t-test). **C.** Er81 protein levels in CINs of the striatum at P6 (n = 12 mice) and P30 (n = 10 mice). Data normalised to the background of the images. **D.** ChAT^+^ interneurons in P30 striatum of a control (*ChAT-Cre; Er81^+/+^*, left) and an *Er81* cKO (*ChAT-Cre; Er81^f/f^*, right) mouse. Scale bar: 25µm. **E-F.** CIN density in the dorsal striatum at P2 (**E**) and P30 (**F**) in the *Er81* cKO compared to the control conditions (P2: n = 6 mice per group, p = 0.871; P30: n = 8 mice per group, p = 0.539). **G.** Lhx6 protein expression in the control (black, n = 5 mice) and the *Er81* cKO conditions (red, n = 5 mice, p = 0.336). **H.** Expression of key molecular markers of cholinergic cells in control and *Er81* cKO conditions (n = 3-5 mice for each group). *Lhx6*; LIM homeobox 6; p = 0.843, *ChAT*: choline acetyltransferase, *Isl1*: Islet1; p = 0.932, *slc5a7*: choline transporter. Data are presented as mean ± SEM. **\*** p < 0.05, **\*\*\*** p < 0.001.

### Cholinergic interneuron properties change in the absence of Er81

To determine the role of Er81 in CIN specification and maturation, we assessed the molecular properties of these cells following Er81 deletion. The LIM homeodomain transcription factor *Lhx6* is expressed in about half of striatal CINs (Lozovaya et al., 2018)and is necessary for MGE-derived GABAergic interneuron specification (Liodis et al., 2007) (Fragkouli et al., 2009) (Flandin et al., 2011). We first examined whether the expression of Lhx6 and Er81 were correlated in the striatal CINs. Er81 protein was similarly expressed in the *Lhx6*-positive and *Lhx6-*negative CINs (**Figures S1C**&**D**), suggesting that *Er81* and *Lhx6* expressions segregate different subtypes of CINs. Consistently, the percentage of Lhx6 protein-expressing CINs (data not shown) as well as the Lhx6 protein and mRNA levels were not different between the control and the *Er81* cKO CINs (**Figure 1G&H**). While we could not detect any Lhx6 protein expression at P6 in control and Er81 cKO mice, we found that the *Lhx6* mRNA levels in CINs were elevated in the absence of Er81 (**Figure S2A**, middle). Moreover, analysis of GABAergic markers showed no changes in glutamic acid decarboxylase GAD65/67 protein (data not shown) and *GAD2* mRNA levels in the striatum (**Figure S2A**, left) in the absence of *Er81*. These results suggest that *Er81* deletion does not affect the GABAergic cell identity of the CINs. We then quantified the expression of genes associated with striatal cholinergic cell identity by fluorescence-activated cell sorting (FACS) followed by quantitative PCR. The expression of the LIM homeodomain transcription factor Islet1 (*Isl1*) (Allaway and Machold, 2017) was significantly increased in the *Er81* cKO at P6 (**Figure S2A**, right), without any change in LIM homeodomain transcription factor *Lhx7 (Lopes et al., 2012)*, *Zic4* (Magno et al., 2017), acetylcholine esterase *AChE,* or choline transporter *Slc5A*7 mRNA levels (**Figure S2A**, left). However, at P30, *Isl1* mRNA levels were not altered, whereas *Slc5A7* expression was upregulated in the absence of *Er81* (**Figure 1H**). Choline acetyltransferase (*ChAT*) mRNA and protein expressions also significantly increased in the *Er81* cKO at P6 (**Figure S2A**). On the other hand, *ChAT* mRNA levels were lower in the *Er81* cKO conditions at P30 (**Figure 1H**), with no change in ChAT protein levels (**Figure S2A**, right), which might be indicative of early maturation of the cells. As Er81 incites a number of molecular changes during developmental stages, the morphology of CINs could be altered when it is removed (Abe et al., 2011). Therefore, we analysed the CIN dendritic properties and ChAT^+^ neuropil. Whilst the overall spread of the dendritic field was unchanged (**Figure 2A&B**), the volume, number of branch points, and Sholl intersections showed a significant reduction in CIN dendritic complexity in the absence of Er81 (**Figure 2C-E**). In addition, we analysed CIN axonal projections and found that ChAT^+^ neuropil area was significantly decreased in the *Er81* cKO condition (**Figure 2F**). Together, these results show that Er81 regulates key developmental genes and the morphological complexity of the cholinergic interneurons of the striatum.

**Figure 2:**
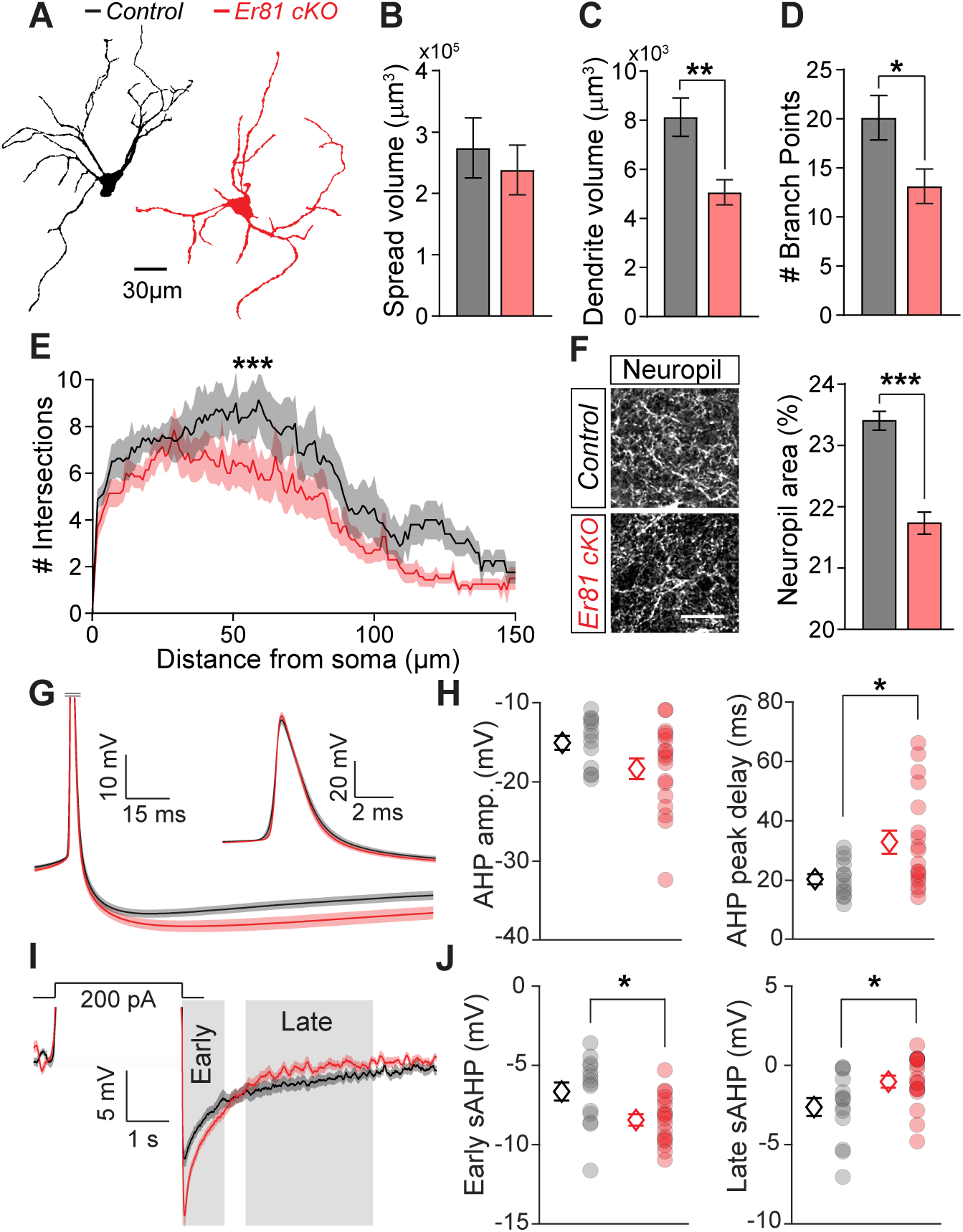
Cholinergic cell properties are altered in the absence of *Er81*. **A.** Morphological reconstruction of soma and dendrites of control and cKO CINs at P30. **B-D.** Volume of dendritic spread (**B**; p = 0.591), total dendritic volume (**C**) and the number of the dendritic branch points (**D**) in the control (n = 8 cells, 3 mice) and the *Er81* cKO CINs (n = 7 cells, 3 mice). **E.** Sholl analysis representing the complexity of the dendritic field in the control (n = 8 cells, 3 mice) and the *Er81* cKO (n = 7 cells, 3 mice). **F.** Quantification of ChAT^+^ neuropil in the striatum (control; n = 4, *Er81* cKO; n = 3 mice, scale bar: 25µm) **G.** Average spike waveforms (± SEM) in the control (n = 14 cells, 4 mice) and the *Er81* cKO (n = 18 cells, 2 mice). Note that the waveform is truncated and rescaled as inset. **H.** AHP properties in the control and the *Er81* cKO mice. Left: AHP amplitude (p = 0.063). Right: AHP decay (p = 0.015, sample size as in G). **I.** Average membrane potential after AP truncation. Grey shadings mark early and late phases of the slow AHP (sAHP, sample size as in G). **J.** Average of the early (left) and the late phase of sAHP (right) in the two conditions (sample size as in G). Data are presented as mean ± SEM. **\*** p < 0.05, **\*\*** p < 0.01, **\*\*\*** p < 0.001.

### Cholinergic interneurons display enhanced intrinsic currents after Er81 deletion

Alterations in molecular and morphological features of CINs in the *Er81* cKO mice suggest that the Er81 transcription factor could also regulate some functional properties of these cells. To address that, we performed *in vitro* whole-cell patch clamp recordings from GFP-labelled ChAT-positive (ChAT^+^) neurons in control (*ChAT-cre; RCE-GFP; Er81^+/+^*) and *Er81* cKO mice (*ChAT-Cre; RCE-GFP; Er81^f/f^* mice). Whilst we could not find any correlation between the expression of Er81 and most of the CIN basic membrane properties at P6 and P30 in control conditions (Table 1), we did observe changes after Er81 deletion from CINs at P6 (Table 2) and P30 (Table 3). In particular, the afterhyperpolarisation (AHP) phase of the action potentials was altered in the *Er81* cKO condition at P30, with significantly slower AHP kinetics compared to the control (Figure 2G&H, Table 3). The absence of Er81 also led to significant changes in both early and late components of the induced slow afterhyperpolarisation (sAHP). While the early component amplitude was larger, the late component amplitude was smaller in the *Er81* cKO, compared to the control CINs (**Figure 2I&J** and Table 3), and we found no change in spike recovery time (Table 3: AP recovery time). This suggests that the overall duration of the pause in CIN firing is unlikely to be affected by *Er81* deletion.

To further analyse CIN physiological properties, we recorded the evoked firing responses and the tail currents responsible for sAHP (Wilson and Goldberg, 2006). CINs exhibited less persistent evoked firing in the *Er81* cKO mice (**Figure 3A**). While the control CINs maintained a high rate of activity during stimulation, the instantaneous firing rate of the *Er81* cKO cells quickly dropped (**Figure 3B**). The average firing rate of the CINs was consistently lower in the *Er81* cKO conditions for all current intensities (**Figure 3C**). Together, our results indicate that the initial responsiveness of the CINs is not altered, but their overall excitability is reduced due to stronger firing rate adaptations in the absence of *Er81* (**Figure 3D**). It has been shown that CIN firing rate is controlled by the large, calcium-dependent delayed rectifier current I_Kr_ (Zhang et al., 2018) (Wilson and Goldberg, 2006). Therefore, the observed differences (**Figure 3B**) might be due to an increase in the I_Kr_ current in the *Er81* cKO conditions. To test this, we applied a depolarising pulse in voltage-clamp mode to activate the outward I_Kr_ current (Wilson and Goldberg, 2006), which was maintained for few seconds after the voltage pulse (**Figure 3E**). We observed that the amplitude of I_Kr_ current was significantly larger in the *Er81* cKO compared to the control CINs (**Figure 3F&G**).

**Figure 3:**
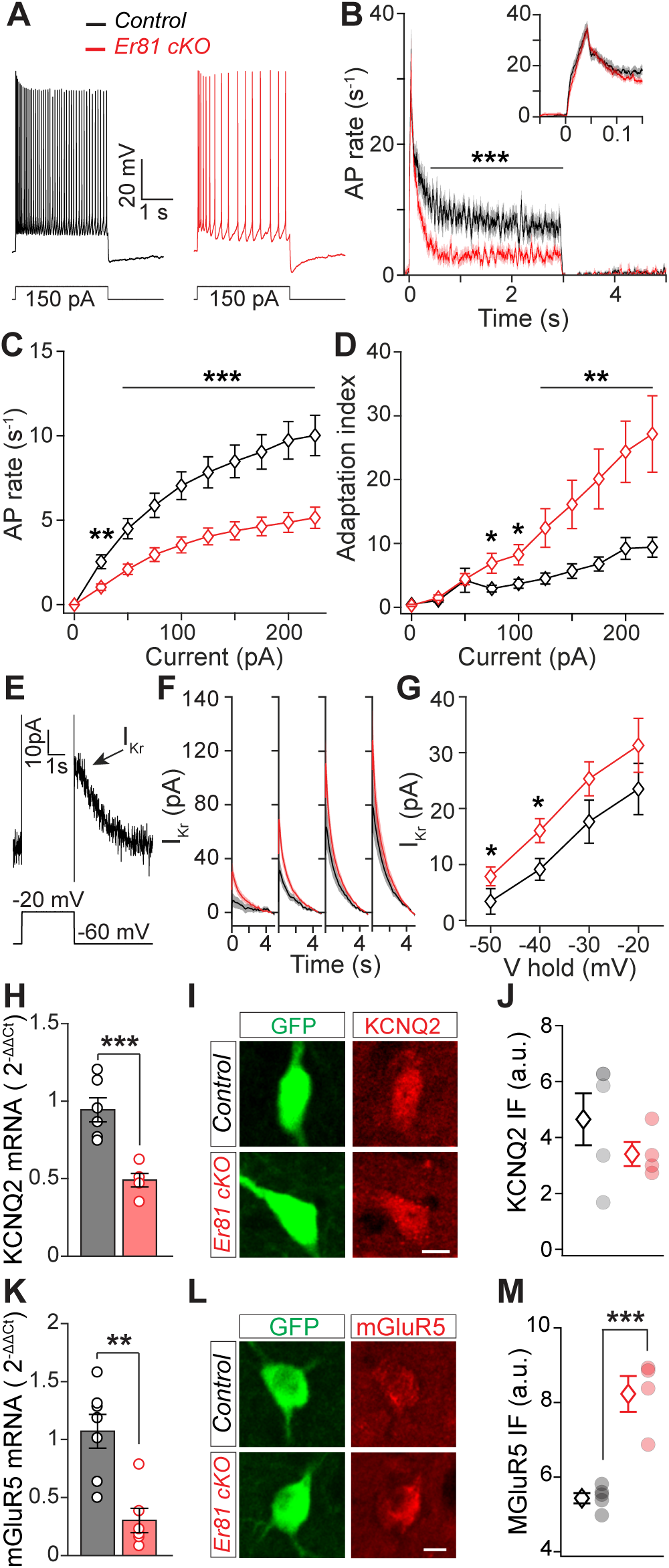
*Er81* deletion reduces cholinergic interneuron excitability due to enhanced delayed rectifier current I_Kr_. **A.** Example responses of the CINs to a positive current pulse in the control and the *Er81* cKO conditions. **B.** Average action potential rate during across all positive current pulses. The inset shows a time-expanded view of the early response (0-0.1 s; p = 0.280, n = 16 cells from 5 control mice and 18 cells from 2 *Er81* cKO mice). **C.** Average action potential rate during positive current steps (sample size as in B). **D.** Adaptation index as a function of the current step amplitude (sample size as in B). **E.** Example trace of the putative I_Kr_ following a depolarising voltage pulse. **F.** Mean traces of I_Kr_ as a function of time at different holding potentials (-50 to −20 mV, 10 mV steps). **G.** Average I_Kr_ as a function of the holding potential (control; n = 14, 3 mice and cKO; n = 16, 2 mice, -30 mV; p = 0.138, −20 mV; p = 0.272). **H-J**. Expression of KCNQ2 mRNA (**H**, n = 6 control and n = 5 *Er81* cKO), examples of CINs (**I**, green) stained for KCNQ2 protein (**I**, red) and the summary of the protein expression (**J**, n = 5 control mice and n = 4 *Er81* cKO mice, p = 0.302). **K-M.** Expression of mGluR5 mRNA (**K**, n = 7 control and n = 6 *Er81* cKO), examples of CINs (**L**, green) stained for mGluR5 protein (**L**, red) and the summary of the protein expression (**M**, n = 5 control mice and n = 4 *Er81* cKO mice). Data are presented as means ± SEM. **\*** p < 0.05, **\*\*** p < 0.01, **\*\*\*** p < 0.001.

In order to decipher the molecular mechanisms behind this regulation, we first analysed the expression of previously identified I_Kr_ mediators in CINs, the KCNQ2/3 hetero-tetramer channels (Zhang et al., 2018). We found a significant decrease in *KCNQ2* mRNA levels (**Figure 3H**), without any change in KCNQ2 protein levels (**Figure 3I** & **J**) between the P30 control and *Er81* cKO mice. We then measured the expression of the metabotropic glutamate receptor 5 (mGluR5), as it can amplify I_Kr_ by raising intracellular calcium concentration (Reiner and Levitz, 2018). Interestingly, we found a decrease in the expression of *GRM5* mRNA (**Figure 3K**) and an increase in mGluR5 protein levels in Er81 cKO conditions (**Figure 3L**&**M**), demonstrating that Er81 deletion alters mGluR5 expression and CIN intrinsic properties at P30. Since *Er81* deletion did not cause any significant change in AHPs, I_Kr_, or their associated properties at P6 (**Figure S2B-H**), we conclude that the ablation of the Er81 expression has an impact on cell maturation, and potentially leads to permanent changes in cell properties in mature state.

A large inward rectifier current is associated with a longer pause in firing and a reduction in CIN spontaneous activity, as it lasts for several seconds (Wilson and Goldberg, 2006).

However, despite a larger I_Kr_ current (**Figure 3F**), we saw no difference in the action potential recovery time (i.e duration of the pause; **Table 2**) and baseline firing rates (**Table 2**). Moreover, the late sAHP showed lower amplitude (**Figure 2I**), which imply that a counter current might be engaged to balance CIN activity in the absence of *Er81*.

The prominent inward rectifying I_h_ current contributes to a rebound activity following hyperpolarisation and the tonic firing of the CINs (Bennett et al., 2000). To examine whether the I_h_ current is also modified in the absence of Er81, we delivered hyperpolarising current pulses to control and *Er81* cKO CINs and found that the sag ratio and I_h_ amplitude remained unchanged between the two conditions at P6 (**Figure S2I-K**). However, both the sag ratio and rebound firing were enhanced at P30 in the absence of the Er81 (**Figure 4A-C**). Consistently, we also found significantly larger I_h_ current amplitude in the *Er81* cKO compared to the control CINs (**Figure 4D-F**). The hyperpolarization activated cyclic nucleotide gated potassium and sodium channel 1 and 2 (HCN1 and HCN2) are the main channel subunits underlying the I_h_ current in the CINs (Zhao et al., 2016). Therefore, we examined whether a potential increase in the expression of these proteins contributes to the enhanced I_h_ current we observed in the Er81 cKO condition. However, we did not find any difference in the expression of neither HCN2 nor HCN1 between the *Er81* cKO and the control CINs (**Figure 4** **G**-**J**). These results suggest that rather than a quantitative change in the channel subunits underlying I_h_, their function is likely enhanced in the absence of Er81. Altogether our results suggest that the absence of *Er81* affects the two major intrinsic currents I_Kr_ and I_h_ in the CINs, potentially through a process involving an upregulation of mGluR5 and its signalling pathways.

**Figure 4:**
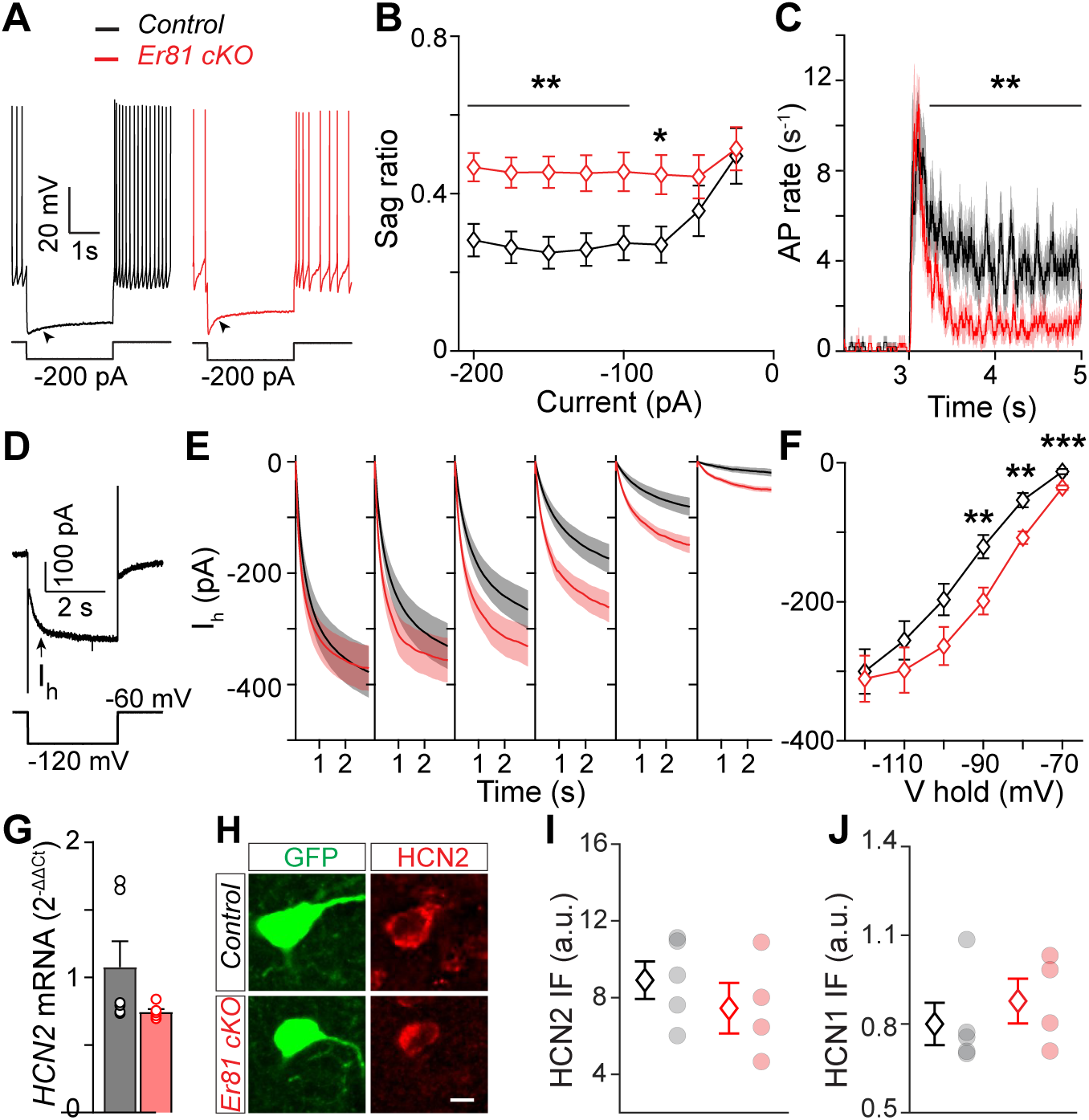
*Er81* deletion amplifies the inward rectifier current (I_h_) in P30 cholinergic interneurons. **A.** Response of a control and an *Er81* cKO CINs to a negative current pulse. Arrow heads show the voltage sag. **B.** Average sag ratio at different current steps (control; n = 16 cells, 5 mice, *ER81* cKO; n = 18 cells, 2 mice, −50 pA; p = 0.133, −25 pA; p = 0.780). **C.** Rebound firing at the end of negative current steps (sample size as in B, average of 3-3.23 s, p = 0.593) **D.** An example voltage-clamp trace shows the slowly activated I_h_ (arrow) during a hyperpolarizing voltage pulse. **E.** Traces of I_h_ averaged across CINs at different holding potentials (-120 to −70 mV, 10 mV steps, control; n = 18 cells, 4 mice, *Er81* cKO; n = 14 cells, 2 mice). **F.** Time averaged I_h_ across CINs at different holding potentials (sample size as in E, −120 mV; p = 0.829, −110 mV; p = 0.340 and −100 mV; p = 0.077). **G-I.** Expression of HCN2 mRNA in the control and the *Er81* cKO mice (**G**, n = 6 mice per group, p = 0.110), example of CINs expressing HCN2 protein (**H,** scale bar is 10 µm) and average HCN2 protein expression in the control (n = 5 mice) and the *Er81* cKO group (n = 4 mice, p = 0.391). **J.** Expression of HCN1 protein in the control (n = 5 mice) and the *Er81* cKO group (n = 4 mice, p = 0.730). Data are presented as means ± SEM. **\*** p < 0.05, **\*\*** p < 0.01, **\*\*\*** p < 0.001.

### Synaptic properties of the cholinergic interneurons are altered in the absence of Er81

Changes in the functional properties of CINs is likely to affect their synaptic integration within the striatal microcircuit. To test whether the deletion of *Er81* alters CIN synaptic profile, we performed *in vitro* patch-clamp recordings of the excitatory and inhibitory spontaneous postsynaptic currents (sEPSCs and sIPSCs) onto CINs. We did not observe any significant changes in the frequency and amplitude of sEPSCs in the absence of *Er81* at P6 or P30 (**Figures S3A&B and 5A-C**). However, the sEPSC decay time was reduced in the *Er81* cKO mice at P30 (**Figure 5D**) but not at P6 (**Figure S5C**), suggesting a potential fine-tuning of the kinetics of the postsynaptic α-amino-3-hydroxy-5-methyl-4-isoxazolepropionic acid (AMPA) and kainate receptors. As CINs receive excitatory inputs from both the cortex and thalamus (Ding et al., 2010), we examined whether pathway-specific inputs were modified in the absence of *Er81*. Bouton analyses revealed no significant changes in the density of the cortical vesicular glutamate transporter type 1 (VGluT1) and thalamic vesicular glutamate transporter type 2 (VGluT2) positive boutons (Doig et al., 2014) on CINs in the absence of *Er81* at either P6 (**Figure S5G**&**H**) or P30 (**Figure 5E-H**). These results suggest that higher mGluR5 protein levels observed in the absence of Er81 is not an adaptive response to changes in glutamatergic synaptic inputs onto CINs.

**Figure 5:**
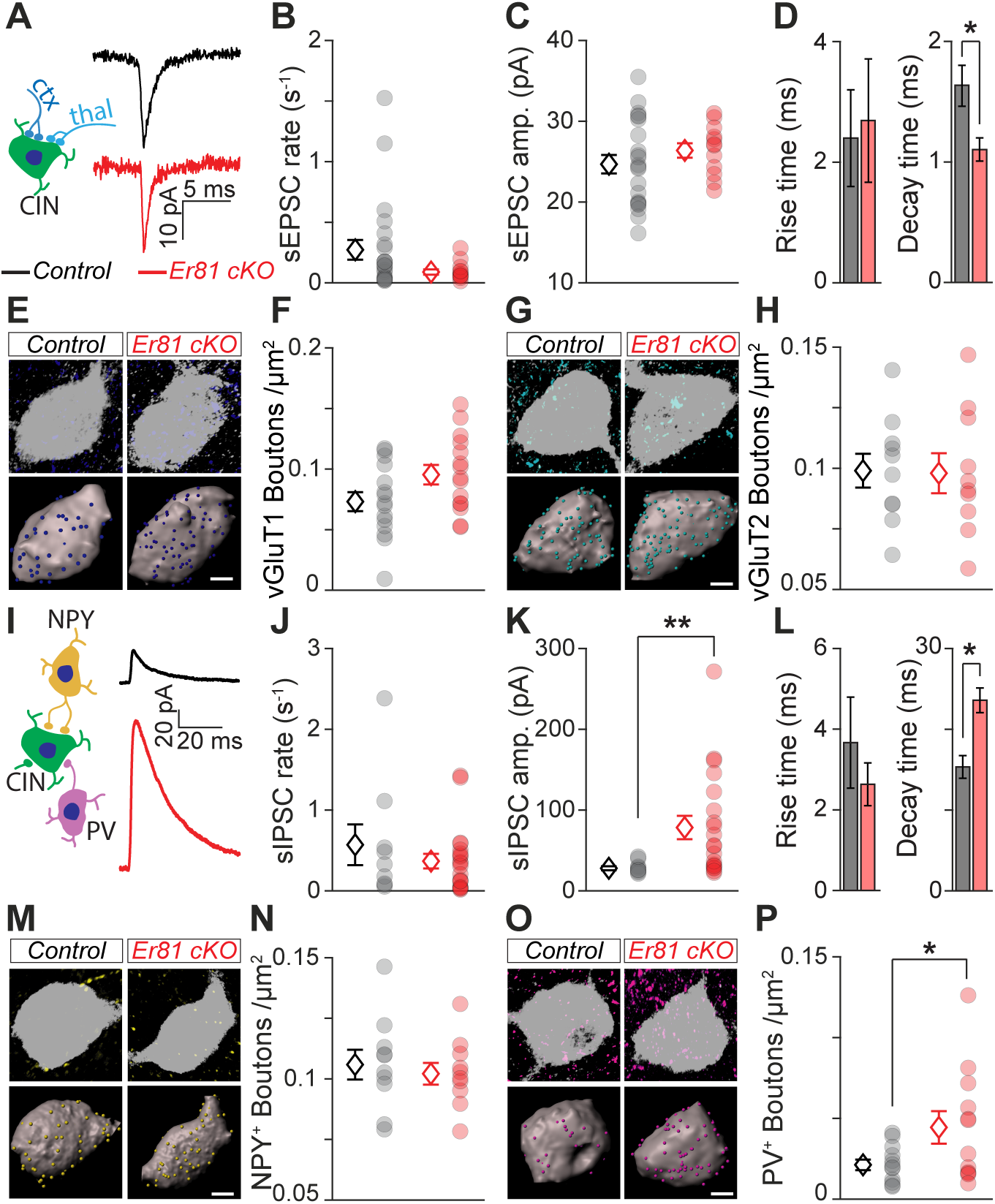
Synaptic properties of the cholinergic interneurons are altered in the absence of *Er81*. **A.** Schematic of excitatory inputs to CINs from cortex (ctx, dark blue) and thalamus (thal, light blue). Average spontaneous excitatory postsynaptic currents (sEPSC) traces of a control and an *Er81* cKO CIN. **B-D.** Rate (**B**, p = 0.188), amplitude (**C**, p = 0.317) rise time (**D**, left, p = 0.871) and decay time (**D,** right) of sEPSCs across CINs (control; n = 22 cells, 9 mice, *Er81* cKO; n = 12, 2 mice). **E.** Top; example CINs (grey) stained for vesicular glutamate transporter 1 (vGluT1, dark blue) representing cortical boutons. Bottom; 3D reconstruction of soma (grey) and vGluT1 boutons (blue spots). **F.** Quantification of vGluT1 bouton density (control, n = 15 cells, 3 mice, *Er81* cKO, n = 15 cells, 3 mice, p = 0.058). **G.** Example CINs stained for PV^+^ boutons (light blue). Details as in E. **H.** Quantification of vGluT2 bouton density (control, n = 10 cells, 3 mice, *Er81* cKO, n = 10 cells, 3 mice, p = 0.392). **I.** Schematic of local inhibitory inputs to CINs from NPY^+^ (yellow) and PV^+^ (magenta) interneurons. Average spontaneous inhibitory postsynaptic currents (sIPSC) traces of a control and an *Er81* cKO CIN. **J-L.** Rate (**J**, p = 0.912), amplitude (**K**) rise time (**L**, left, p = 0.391) and decay time (**L,** right) of sIPSCs across CINs (control; n = 10, 2 mice, cKO; n = 20 cells, 3 mice). **M.** Example CINs stained for NPY^+^ boutons (yellow). Details as in E **N.** Quantification of NPY^+^ bouton density (control, n = 10 cells, 2 mice, *Er81* cKO, n = 10 cells, 2 mice, p = 0.627). **O.** Example CINs stained for PV^+^ boutons (magenta). Details as in E. **P.** Quantification of PV^+^ bouton density (control, n = 12 cells, 3 mice, *Er81* cKO, n = 12 cells, 3 mice). Scale bars is 5 µm. Data are presented as means ± SEM. **\*** p < 0.05, **\*\*** p < 0.01. Averages are shown as diamonds and individual neurons are plotted as circles.

We then quantified inhibitory inputs onto the CINs. Recordings of sIPSCs showed similar frequency (**Figure 5I&J**) but significantly larger amplitude in the *Er81* cKO compared to the control cells at P30 (**Figure 5K**), with no changes at P6 (**Figure S3D-F)**. The decay time of the sIPSCs was also larger in the *Er81* cKO compared to controls (**Figure 5L**). Since interneurons expressing the neuropeptide Y (NPY^+^) are strongly connected to CINs (Straub et al., 2016, English et al., 2011), we determined whether NPY^+^ bouton density onto CINs are altered in the absence of Er81. We found no difference between the two groups at P30 (**Figure 5M&N**). However, we observed a higher density of NPY^+^ boutons in the *Er81* cKO compared to control mice at P6 (**Figure S3I**). PV^+^ boutons were denser on the *Er81* cKO CINs compared to controls at P30, with no change in NPY^+^ boutons (**Figure 5O-P**). Overall these results suggest that the increase in IPSC amplitude of mature CINs lacking Er81 is due to a stronger connection with PV interneurons.

As *Er81* deletion affects cholinergic neuropil (**Figure 2F**), we expected that CIN output to other striatal cell types would also be altered. We found that the density of boutons expressing the specific CIN synaptic terminal marker vGluT3 (Higley et al., Nelson et al.) was increased at P30 but not at P6 on neighbour CINs in the absence of Er81 (**Figure S4A&B**). On the contrary, vGluT3 bouton density onto NPY^+^ interneurons was reduced at P6 but not at P30 (**Figure S4C**) and was significantly reduced onto P30 PV-expressing interneurons in cKO conditions (**Figure S4D**). These results show that the specific ablation of *Er81* affects CIN connectivity with other interneurons of the striatum. Together with enhanced intrinsic currents, these changes in connectivity are likely to alter the whole *in vivo* striatal activity.

### Er81 deletion enhances responsiveness of striatal units in awake mice

To determine the consequences of the alterations observed *in vitro*, we performed multi-electrode array recordings from the dorsal striatum coupled to whisker stimulation in awake mice (**Figure 6A&B**). We recorded a total of 2087 units from 3 control and 4 *Er81* cKO mice. Putative fast spiking cells characterized by narrow spikes (Mallet et al., 2005) (Lee et al., 2017) were excluded and the remaining units were sorted based on their average evoked firing rate.

**Figure 6:**
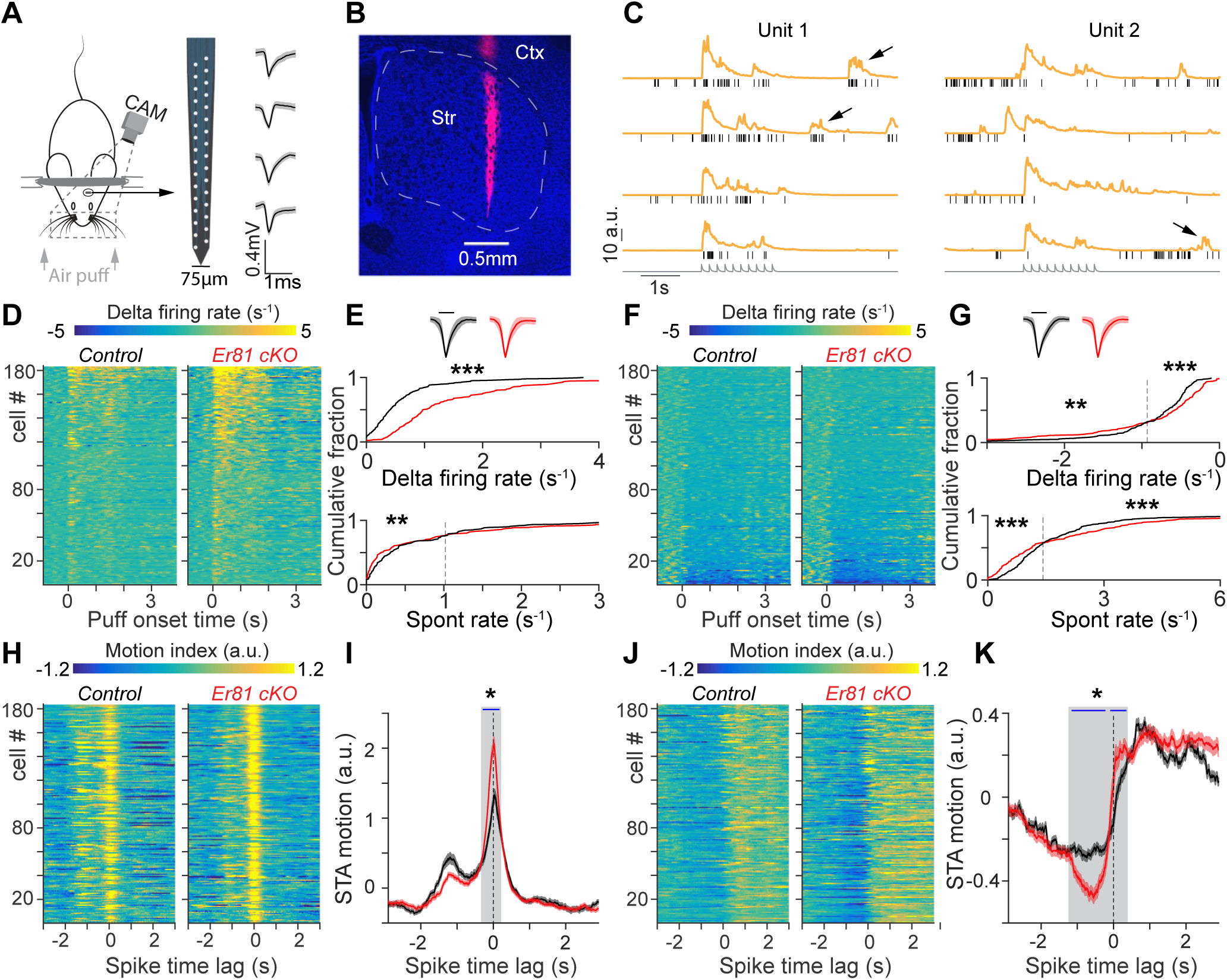
Enhanced responsivity of striatal units in the absence of cholinergic *Er81*. **A.** Head-fixed awake recording by multi-electrode arrays (middle) is depicted. Right: example traces of sorted spikes. **B.** Track of the recording array is marked by DiI (magenta). Dashed area outlines dorsal striatum. **C.** Example trials (4 out of 20 trials per unit) showing spike rasters (black) and the combined movements of the whiskers and the nose quantified as motion index for 2 units (orange). Grey traces at the bottom show the air puff trains. **D.** Normalised spiking activity of the excited cells (control; n = 183, 3 mice, *Er81* cKO; n = 183, 4 mice). **E.**: Spike waveforms (top; scale bar; 500 ms), cumulative distribution of the response magnitude (middle) and cumulative distribution of spontaneous spike rates (bottom) of the excited units (n = 183 per group; control in black and *Er81* cKO in red). Dashed line shows the point of intersection of the estimated cumulative functions (right portion: p = 0.098). **F.** Colour plots represent normalised and sorted spiking activity of supressed cells (control: n = 183, 3 mice; *Er81* cKO; n = 184, 4 mice). **G.** Spike waveforms (top; scale bar; 500 ms), cumulative distribution of the response magnitude (middle) and cumulative distribution of spontaneous spike rates (bottom) of the inhibited units. **H.** Spike triggered average (STA) of the motion index corresponding to the excited units (same units as in D). **I.** Average STA across the excited units (same units as in D and H). Dashed line shows the spike time and blue line shows significant time points within the shaded region (two-sided permutation test). **J.** STA of the motion index corresponding to the inhibited units (as in F). **K.** Average STA over the inhibited units (as in F and J). Dashed line shows the spike time and blue lines show significant time points within the shaded region (two-sided permutation test). Traces are presented as means ± SEM. **\*** p < 0.05, **\*\*** p < 0.01, **\*\*\*** p < 0.001.

Striatal projection neurons have low spontaneous activity and show phasic responses to sensorimotor stimuli (Inokawa et al., 2010) while CINs exhibit a suppression or pause of their ongoing tonic activity during cortical and thalamic inputs (Ding et al., 2010) {Doig, 2014 #4427. Therefore, we analysed cells that are respectively activated or depressed during whisker stimulation (referred as ‘excited’ and ‘inhibited’ units, respectively; **Figure 6C**). In the ‘excited cell’ population (**Figure 6C**, left), we found significantly larger responses to the stimulus in the *Er81* cKO mice compared to the control (**Figure 6D&E** top and **Figure S5E**). In this group, putative output neurons characterized by a low spontaneous firing rate (i.e. < 1 Hz; see methods), were found to fire at lower rates in the *Er81* cKO condition (**Figure 6E**, bottom). In the ‘inhibited cell’ population, representing the putative CINs (**Figure 6C** **right; Figure S5H)**, the most inhibited units (with responses < -0.93 s^-1^), showed a stronger response to the stimulus in the *Er81* cKO compared to the control mice (Figure 6F&G top). On the other hand, stimulus-induced suppression of the cells experiencing less inhibition (> -0.93 s^-1^) was reduced in the *Er81* cKO compared to the control mice (Figure 6G top). Regarding the spontaneous activity of the inhibited cells, we observed a higher activity of the cells firing at a rate above 1.42 s^-1^ and a lower activity of the cells firing below 1.42 s^-1^ in the *Er81* cKO condition compared to control (Figure 6G, bottom). These results show that more putative CINs are recruited (i.e. prominent pause and rebound) by the stimulus in the absence of *Er81*.

Striatal cell firing could be affected as head-fixed awake mice voluntarily move {Ranjbar-Slamloo, 2019 #4285}. We thus analysed how snout and whiskers behaved during the course of stimulation. We found that the cumulative distribution of the motion index was temporally different, revealing a higher motor activity of the *Er81* cKO mice compared to control (**Figure S5A-C**). We then performed a reverse correlation analysis to examine the coupling between spikes and movements. The spike-triggered average (STA) motions showed a larger peak of motion at the spike time in the ‘excited’ cell group (i.e. the putative output neurons) of the *Er81* cKO mice compared to control (**Figure 6H**&**I**). Magnitude of the motions near spike time was also larger in the ‘suppressed’ cells (**Figure 6J&K**), indicating a better temporal alignment of the spikes to the onset of the movement in the *Er81* cKO mice. Negative to positive STA near the spike (**Figure 6K**) confirms that most of the cells were suppressed following the sharp transitions from quiet to active states (**Figure 6C**, right). Overall, this analysis shows that striatal neurons are more responsive to the sensorimotor inputs. To examine the relative contributions of the stimulus and the motion in generating spikes in the absence of Er81, we performed cross-correlation analysis between the firing rate, motion, and puff stimulation. In the ‘excited cells’ (i.e putative output neurons), positive spike-motion correlations were on average stronger in the absence of *Er81*, while the corresponding spike-puff correlations were not affected (**Figure S5D-F**). On the other hand, negative correlations in the ‘inhibited cells’ (i.e putative CINs) were significantly larger in the *Er81* cKO compare to the control mice **(Figure S5G-I**). Interestingly, we also found that spike-motion correlations were consistently higher than spike-puff correlations (**Figure S5F&I**), suggesting that neurons were mostly tuned to the voluntary movements, rather than the stimulus. Altogether our results indicate that physiological changes in the *Er81* cKO mice lead to higher motor activity and a more effective modulation of unit activity in the striatum. These data imply that the putative CIN population producing phasic responses during movement are more responsive to sensorimotor inputs in the absence of *Er81,* and cause a higher excitation of the putative output neurons.

### Er81 deletion from cholinergic cells alters habit formation

Striatal CINs are crucial for cognitive flexibility (Prado et al., 2017) (Okada et al., 2014) (Okada et al., 2018). To assess the impact of *Er81* deletion on goal-directed behaviours, we designed a reversal learning task, allowing the assessment of cognitive flexibility and habit formation. In the first part of the training, both control and *Er81* cKO mice were able to discriminate the rewarded corridor of the maze (**Figure 7A**, left), and switch to the opposite side after a first reversal on Day 4 (**Figure 7A**). *Er81* cKO mice hence displayed intact learning capacities (Days 1 to 3) as well as cognitive flexibility (Days 4 to 8). Mice were then over-trained for 5 days after they reached their performance plateau, in order to induce habit formation (Balleine, 2019) (**Figure 7A**, middle). Interestingly, *Er81* cKO mice performed significantly higher compared to controls during the second reversal (Day 14; **Figure 7A**, right). To further analyse this session, we compared the first five trials to the last five trials, and found that controls remained significantly below chance level throughout the whole session (**Figure 7B**). This inability to flexibly shift to the new rewarded side indicates that control mice were engaged in habitual behaviour. In contrast, *Er81* cKO mice displayed rapid improvement of their performance, reaching chance level by the end of the trials. This result was further confirmed by the following session during which *Er81* cKO mice performed significantly above chance level, unlike controls (**Figure 7A**, right). Together, these results show intact goal-directed actions associated with an impairment of habit formation in mice lacking Er81. These mice also show general hyperactivity as they move faster in the middle corridor (**Figure 7C**). These results are consistent with a higher motion index of the snout and whiskers in the head-fixed experiments (**Figure S5C**).

**Figure 7:**
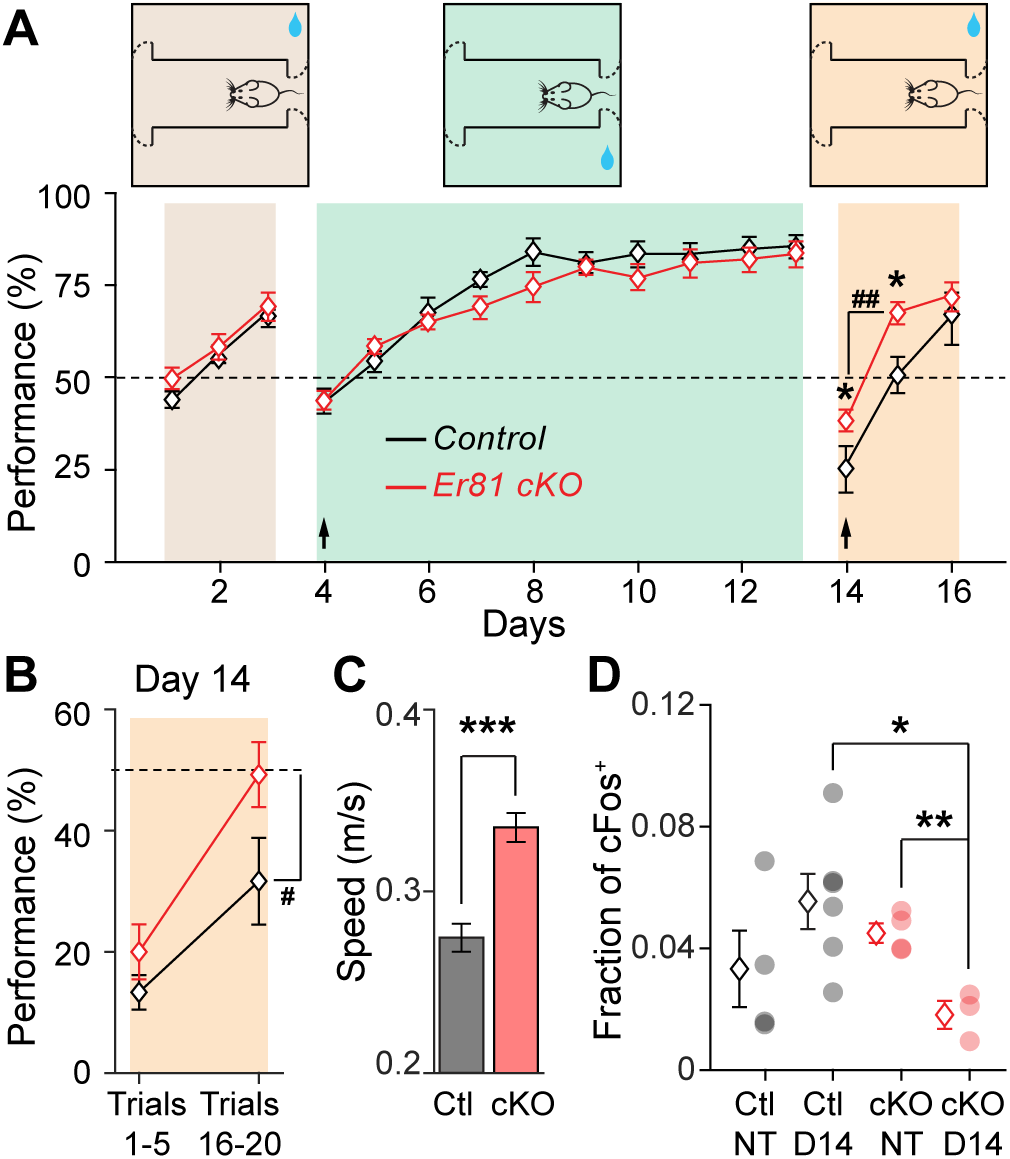
*Er81* cKO mice exhibit disrupted habitual behaviour. **A.**: Mice performed a reversal learning task in a 3 corridors arena (top). Dashed curve: one-way gate. Blue droplet: sucrose reward. Performance of the mice in each session (bottom; control in black vs *Er81* cKO in red, n = 4-13 mice). Arrows indicate the days of reversal. **B.** Average performance across the first 5 trials and last 5 trials of the day 14 (control; n = 12 mice, *Er81* cKO; n = 13 mice). **C.** Average locomotion speed of the mice in the middle corridor (control; n = 124 sessions, 4-11 mice, *Er81* cKO; n = 133 sessions, 4-12 mice). **D.** Fraction of the cells expressing cFos in non-trained (NT) and day 14 of training (D14) in the control (n = 4 NT and n = 6 D14, p = 0.179) and the *Er81* cKO (n = 4 NT and n = 3 D14; interactions between training and genotype: p = 0.022, multiple regression test). Data are presented as means ± SEM. **\*** p < 0.05, **\*\*** p < 0.01, **\*\*\*** p < 0.001 (*Er81* cKO vs the control). **^#^** p < 0.05, **^##^** p < 0.01 (versus the chance level, one-sampled t-test). Dashed lines show the chance level (performance at 50%).

We then analysed the changes in striatal activity associated with the paradigm. Expression of the cellular activity marker cFos in the dorsal striatum revealed an alteration of the task-induced activity of the output neurons in the Er81 cKO mice (**Figure 7D**). While training had no effect upon cFos expression in controls, the *Er81* deletion induced a drastic hypo-activation of the striatum in trained mice, compared to non-trained mice. The activity of the striatum being crucial for habitual actions, this data could explain the inability of the *Er81* cKO mice to form habits, overall resulting in improved cognitive flexibility.

## DISCUSSION

Here, by utilising *in vitro* and *in vivo* assessments, we reveal fundamental alterations in cholinergic cell properties in the specific absence of Er81 expression that render the striatal network more responsive to sensorimotor inputs and lead to disrupted habitual behaviour.

### Control of the cholinergic cell identity

The functional implications of molecular heterogeneity in striatal CINs are at present not well understood (Munoz-Manchado et al., 2018) (Lozovaya et al., 2018) (Magno et al., 2017). CINs originate from distinct areas of the subpallium which give rise to heterogeneous populations of cholinergic cells in the forebrain (Ahmed et al., 2019). These areas are further divided into subdomains, based on the combinatorial expression of multiple transcription factors such as Lhx6, Lhx7, Isl1 and Er81 (Flames et al., 2007) (He et al., 2016). The temporal order of expression of these transcription factors and their interactions are crucial for normal development of cholinergic neurons (Zhao et al., 2003) (Allaway and Machold, 2017) (Ahmed et al., 2019). In this study, we identified Er81 as a molecular controller of key CIN properties and investigated its interaction with other transcriptional factors. Future fate-mapping studies will highlight more specifically the proportion of Er81-expressing striatal CINs that may originate from different sources (i.e. MGE, POa and septum). The role of Er81 in the dopaminergic (Flames and Hobert, 2009) (Cave et al., 2010) and serotoninergic cell fates (Lloret-Fernandez et al., 2018) and here, in cholinergic cell identity, suggest a general regulatory mechanism of Er81 on the emergence of functional neuromodulatory systems in the developing brain.

In the mature cortex, Er81 directly regulates the functional diversity of cortical parvalbumin-expressing (PV) interneurons (Dehorter et al., 2015). Whilst Er81 expression is strong and mostly nuclear in the cortical PV interneurons, it remains weak and cytoplasmic in the CINs of the striatum. This suggests that Er81 works through different mechanisms in the CINs than in the PV interneurons (Dehorter et al., 2015), likely through protein-protein interactions or regulation of translational processes. However, whilst the molecular targets may be different, Er81 seems to have a common role in neuronal function across different cell types, regulating excitability and spike timing (i.e the phasic activity of the striatal CINs and the firing latency in the cortical (Dehorter et al., 2015) and striatal PV interneurons; unpublished data).

CINs are neurochemically complex as they co-release GABA (Saunders et al., 2015) (Granger et al., 2016) and glutamate (Higley et al., 2011) alongside acetylcholine, to provide both potent inhibition (Zucca et al., 2018) or excitation (Tepper et al., 2018) within the striatum. As we found changes in the molecular properties of the CINs (acetylcholine transporter, Isl1 and ChAT), co-release with other neurotransmitters might be also affected by Er81 deletion and ultimately alter the neurochemical function of the CINs. Our study therefore emphasises the need to further investigate the molecular factors determining the identity and the functional diversity of striatal CINs.

### Mechanisms regulating tonic and phasic activity of the striatal cholinergic neurons

Intrinsic electrophysiological properties are responsible for maintaining tonic activity of striatal CINs (Bennett et al., 2000). Together with synaptic inputs (Franklin and Frank, 2015) (Klug et al., 2018), they also contribute to the phasic response to stimuli (Zhang et al., 2018). The role of molecular factors in the development of these properties was unknown. For the first time, our findings link the Er81 transcription factor with intrinsic currents and plasticity-related receptors such as mGluR5, and provide a basis to determine which molecular factors contribute to the emergence of the unique properties of striatal CINs. Elevated I_Kr_ and I_h_ currents, together with larger inhibitory inputs on CINs, can explain the sharper firing dynamics and higher correlations of the putative CINs activity with sensorimotor inputs *in vivo* in the absence of Er81 (Bennett et al., 2000) (Wilson and Goldberg, 2006) (Zhang et al., 2018). KCNQ2/3 and HCN2 channels in CINs underlie the I_Kr_ and the I_h_, respectively. Whilst our results show no difference in their expression in the absence of Er81, these voltage-sensitive channels could be modulated by second messengers such as phosphatidylinositol-4,5-bisphosphate (PIP_2_) and calcium for KCNQ2/3 (Wilson and Goldberg, 2006) (Falkenburger et al., 2010) (Kim et al., 2016), and cAMP for HCN2 (Zhao et al., 2016) (Alvarez-Baron et al., 2018). We propose that the enhanced I_Kr_ in the *Er81* cKO cells is due to a higher intracellular calcium concentration mediated through mGluR5 (Niswender and Conn, 2010). We also suggest that enhanced cAMP signalling leads to the increase in the I_h_ current. Lower CIN excitability in the absence of *Er81* can indeed result in a decreased acetylcholine recruitment at the CIN membrane, reduced M2 muscarinic receptor activity and consequently, to higher levels of intracellular cAMP (Zhao et al., 2016) (Alvarez-Baron et al., 2018).

We propose that the changes in CIN activity in the absence of Er81 modifies their connectivity and morphology via homeostatic compensations. However, we cannot confirm the origin or the circuit-level functional consequences of the synaptic changes. As CINs control the striatal output neurons (Mamaligas and Ford, 2016) and modulate their GABAergic (English et al., 2011), glutamatergic (Mamaligas et al., 2019) and dopaminergic inputs (Brimblecombe et al., 2018) (Threlfell et al., 2012) (Kosillo et al., 2016), we expect that alteration in CIN functional properties will have a crucial impact on the striatal sensorimotor processing *in vivo*.

### Implications for sensorimotor processing in the striatum

Our study reveals for the first time that developmental alterations of striatal CINs strongly impact the process of integration of sensorimotor information, which is critical for the acquisition and the update of adaptive actions (Markowitz et al., 2018) (Robbe, 2018). It also highlights the significance of cholinergic signalling for movement control and learning. Putative striatal CINs better encode motion and display enhanced phasic responses in the *Er81* cKO mice. In particular, at the onset of movement, spikes are more time-locked to the events and less jittered. This also enhanced the timing of putative output neurons which directly encode CIN firing (Mamaligas and Ford, 2016). Modifications in the pause of the CINs will therefore strongly affect basal ganglia output and thus information processing during reinforcement learning. Enhanced timing of the striatal units indicates that spike timing-dependent plasticity (STDP) could be affected by the Er81 deletion (Cui et al., 2018), via a metabotropic glutamate receptor-induced LTD or a NMDA receptor-induced LTP (Fino et al., 2008). However, the mechanisms of STDP have been poorly investigated in the CINs as of yet (Perrin and Venance, 2019). More investigation is therefore required to better understand how CIN alterations impact intrinsic, synaptic and structural plasticity and overall, the striatal function.

### Behavioural significance of altering the pause of striatal cholinergic interneurons

CINs control habit formation via the synchronisation and strengthening of the activity of the striatal output neurons (Gritton et al., 2019) (O’Hare et al., 2016) (Yin et al., 2004) (Balleine, 2019). Strong pauses have been shown to emerge during learning in response to reward-associated stimuli but not to neutral stimuli (Graybiel et al., 1994) (Morris et al., 2004). Our study reveals a notable increase in the magnitude of CIN pause responses following the ablation of the Er81 transcription factor. As cells overreact to non-rewarding stimuli (i.e. whisker stimulation), it suggests that the increased strength of the pause is an abnormal condition. Adaptable CIN responses to sensorimotor inputs are required for suppression of competing actions and the expression of habitual behaviour, a key adaptive behaviour that improves the efficacy of actions when engaged in repetitive behaviours (Prado et al., 2017) (Okada et al., 2014) (Okada et al., 2018). Because of its crucial role in behavioural flexibility, it would be of major interest to study the Er81 transcription factor as a potential target to manipulate set-shifting capacities. Moreover, Er81 could contribute to the pathophysiology and cognitive defects commonly observed in disorders where repetitive behaviours are debilitating (Lewis and Kim, 2009), such as autism (Karvat and Kimchi, 2014), obsessive-compulsive disorder (Xu et al., 2015) and addiction (Graybiel and Grafton, 2015).

## METHODS

### Mice

We generated *Er81^+/+^; ChAT-Cre* (control) and *Er81^flox/flox^*; *ChAT-Cre* (*Er81* conditional knockout, cKO) mice by crossing *Er81^flox/flox^* mice (generous gift by Prof Marin at the MRC, London UK) with *ChAT-Cre* mice (#006410 ChAT-IRES-Cre from The Jackson Laboratory). We also generated *Er81^+/+^; ChAT-Cre; RCE-GFP* (control) and *Er81^flox/flox^*; *ChAT-Cre; RCE-GFP* (*Er81* cKO) mice by crossing *Er81^+/+^; ChAT-Cre or Er81^flox/flox^*; *ChAT-Cre* with *Er81^+/+^*; *RCE-GFP or Er81^flox/flox^*; *RCE-GFP* (kindly supplied by Prof Marin at the MRC, London UK). We used the *Er81^+/+^; Nkx2.1-CreER; RCE-GFP* (control) and *Er81^flox/flox^*; *Nkx2.1-Cre; RCE-GFP* (Er81 *cKO*) mice for the analysis of the CIN morphology. *Lhx6-Cre; RCE-GFP* mice were used to analyse Er81 expression regarding the Lhx6 phenotype of CINs. All experiments were conducted with approval from the Australian National University Animal Experimentation Ethics Committee (protocol numbers A 2016/14, A2018/43 and A2018/66). All efforts were made to minimise suffering and reduce the number of animals. Only male mice were used in this study.

### *In vitro* electrophysiological recordings

We used *Er81^+/+^; ChAT-Cre; RCE-GFP* (control) and *Er81^flox/flox^*; *ChAT-Cre; RCE-GFP* (Er81 mutant) mice at ∼P6 or ∼P30 for slice electrophysiology experiments. Mice were deeply anaesthetised with isoflurane and perfused with ice-cold oxygenated, artificial cerebrospinal fluid (ACSF) containing (in mM): 248 sucrose, 3 KCl, 0.5 CaCl2, 4 MgCl2, 1.25 NaH2PO4, 26 NaHCO3, and 1 glucose, saturated with 95% O2 and 5% CO2. The animals were then decapitated, brain was removed and placed in ice cold oxygenated sucrose-based cutting solution. Sagittal slices of 400 µm were cut using a Leica VT1200S vibratome. Slices were then maintained at room temperature in ACSF containing (in mM): 124 NaCl, 3 KCl, 2 CaCl2, 1 MgCl2, 1.25 NaH2PO4, 26 NaHCO3 and 10 glucose saturated with 95% O2 and 5% CO2. For patch clamp recordings in whole-cell configuration, slices were transferred to a chamber and continuously superfused with ACSF at 34°C. Green fluorescent protein (GFP) expressing cholinergic cells located in the dorsal striatum were visualised by infrared-differential interference optics with a 40x water-immersion objective. For targeting GFP expressing neurons, slices were illuminated by blue light through the objective. Microelectrodes (4-6 MΩ) were pulled from borosilicate glass (1.5 mm outer diameter x 0.86 inner diameter) using a vertical P10 puller (Narishige).

For voltage-clamp recordings, a caesium gluconate-based intracellular solution was used containing (in mM): 120 Cs-gluconate, 13 CsCl, 1 CaCl2, 10 HEPES, and 10 EGTA (pH 7.2-7.4, 275-285 mOsm). We used the caesium-gluconate solution to measure spontaneous and miniature GABA_A_ currents at the reversal potential for glutamatergic (+10 mV) events and glutamatergic currents at -60 mV. For current clamp recordings, we used a potassium-gluconate-based intracellular solution containing (in mM): 140 K-gluconate, 10 HEPES, 2 NaCl, 4 KCl, 4 ATP, and 0.4 GTP. Neurobiotin (2-5 mg/ml) was added for post-recording immunocytochemistry. Electrophysiological signals were low-pass filtered on-line at 10kHz with a Multiclamp 700B (Axon Instruments) amplifier and acquired at a 20-kHz sampling rate with a LIH 8+8 (HEKA) data acquisition board and WinWCP software (created by John Dempster, University of Strathclyde). Circuit capacitance was corrected after gigaseal formation. Series resistance and liquid junction potential were not corrected. Cell attached spikes were recorded for at least 60 seconds, after gigaseal formation in voltage-clamp mode. Following break-in, the test pulse was monitored for a few seconds to ensure a stable, low access resistance (R_a_ < 20MΩ). In whole-cell configuration, spontaneous firing of the CINs were recorded for 60 seconds and then a current pulse of 200pA was delivered for 3 seconds to obtain slow afterhyperpolarization (sAHP). An additional 60 seconds of spontaneous activity was recorded following the pulse. Then, a current-steps protocol (-200 pA to 225 pA, 25 pA steps, 3 s duration and 7 s intervals) was applied to obtain membrane properties and excitability of the cells. In voltage clamp mode, membrane potential was held at -60 mV. Hyperpolarising voltage pulses (-120 mV to −70 mV, 10 mV steps, 3 s duration and 7 s intervals) were applied to measure I_h_. To measure delayed rectifier or I_Kr_ currents, depolarizing voltage pulses were applied (-50 mV to −20 mV, 10 mV steps, 3 s duration and 7 s intervals). To measure postsynaptic currents, the voltage was held at -60 mV for EPSCs or +10 mV for IPSCs. A test pulse was applied every 10 seconds to exclude PSC recordings in case access resistance dropped below 75 percent of the initial value.

Electrophysiological data were analysed in MATLAB using custom written codes. Spontaneous action potential before the first current pulse was taken to examine tonic activity of CINs. Threshold was detected based on the positive peaks occurring above 10 mV/ms in the first derivative of the membrane potential trace. The action potential width was defined as the interval between the first and second threshold crossings. The afterhyperpolarization (AHP) amplitude and its peak delay were obtained at the minimum potential within a 200 ms interval after the second threshold crossing of the AP. A bi-exponential function was fit to the AHP to calculate rise and decay time-constants. To characterise membrane potential dynamics following depolarization, action potentials were removed by linearly interpolating from 30 ms prior to the AP threshold to 100 ms after second threshold crossing point of the AP. The early component of slow APH (sAHP) was calculated by averaging 1 s of the V_m_ following the offset of the current pulse and the late component was averaged from 1.5 s to 4.5 s after the pulse offset. Input resistance (R_in_) was calculated at the minimum of the voltage response (V_min_ to −200 pA pulse, before the sag in V_m_ appeared. Sag ratio was calculate based on the following formula: *Sag ratio = (V_min_ -V_steady_)/ (V_min_ -V_rest_)*, where *V_min_* was the minimum of the voltage during the pulse, *V_steady_* was the average of membrane potential at 2.5-3 second after the onset of the current pulse and *V_rest_* was the medium of the membrane potential within 500 ms preceding the onset of the current pulse. Action potentials were converted to 0 and 1 arrays based on their peak time and then averaged across neurons in 40 µs bins and smoothed with a 40 ms window to obtain instantaneous AP rates (Figure 3B & 4C). AP rate was calculated as the average over the pulse (Figure 3C). Statistical tests on the instantaneous firing rates were applied on intervals were the effect was consistently maintained. Adaptation index was calculated as the range of the inter-spike-intervals divided by their minimum.

In voltage clamp mode, hyperpolarizing and depolarizing voltage pulses were delivered to characterise inward rectifier (I_h_) and delayed rectifier (I_Kr_) currents respectively. To calculate I_Kr_ (tail current), the current at 4-5 second following the offset of the voltage step was subtracted and a 4 second window following the offset was averaged across time. I_h_ was quantified by subtracting the positive peak of the current and then averaging from this peak to up to 2.9 seconds. Postsynaptic currents were isolated using a manual threshold and applying principle component analysis in MATLAB.

### In vivo recordings

*Er81^f/f^; ChAT-Cre* and *Er81^+/+^; ChAT-Cre* mice were used for *in vivo* extracellular array recordings. Animals were kept in individual cages and provided with *ad libitum* food and water. Before the surgical operation, anaesthesia was induced by 3% isoflurane and the head was mounted on a stereotaxic device (Stoelting). Local anaesthetic was applied to the scalp (Lignocaine, Lmx4) and eye gel (Viscotears from Novartis) to both eyes. The skull was exposed and cleared from fascia. A thin layer of tissue adhesive (Vetbond; 3M, St Paul, MN, USA) was applied on the skull. A custom-made head bar was then glued to the skull and secured by dental cement. The cement was applied all over the skull except the area of intended craniotomy which was filled with silicon sealant (Kwik-Kast, WPI, Sarasota, FL, USA) at the end of the surgery. Animals were injected with 5 mg/kg of carprofen and 0.86 ml/kg of penicillin (I.P. injections) and placed on a warm heating pad to recover from anaesthesia. The animals were returned to their home cage and allowed 6 days of recovery from surgery. Then the mice were head-fixed for 3 sessions of 15-90 minutes over 3 days to habituate. On the following day, the mice were anaesthetised with 3% isoflurane and two small craniotomies (< 1 mm in diameter) were drilled (0.0-0.4 mm from bregma, 2.5 mm lateral) on both sides and the dura was left intact and the craniotomy was filled with silicon sealant. The first recording session started at least 3 hours after the recovery from anaesthesia. The recording probe was moved down to a depth of ∼1.2 mm (distance of the tip from the dura) followed by ∼0.1-0.2 mm advancement after each recording. This was repeated up to a depth of ∼4 mm. In the second recording session (24 h later), the opposite craniotomy was used for recording. At the end of second recording session, the mice were perfused transcardially and the brain was removed for histological verification.

To obtain sensory driven responses in striatum, a bilateral train of air puffs (10 pulses of 200 ms/cycle) was applied to the whisker arrays during each recording using a Pico spritzer (Parker Instruments) device. The pressure was adjusted for once to generate visible movements of the whiskers allowing the puffers to be placed in front of the animal, out of the reach of the whiskers. The puff train was repeated every 10 seconds for 20 trials per recording. Electrophysiological data were acquired using 32 channel NeuroNexus double linear arrays (A1x32-Poly2-10mm-50s-177, NeuroNexus, Ann Arbor, MI, USA) coupled with CerePlex Direct data acquisition system (BlackRock Microsystems, Salt Lake City, UT, USA). A high-speed imaging system (Mikrotron, Germany) was used to film the top view of the head at 250 frame/s during electrophysiological acquisition.

The depth of recording at each electrode site was calculated using the probe guidelines (http://neuronexus.com/electrode-array/a1x32-poly2-10mm-50s-177/). The whole span of recording depth for individual channels across animals was 0.37-4.20 mm. Based on histology data, we estimated the boundary between cortex and striatum to be at ∼2 mm. We limited our analysis of striatal units to 2.0-3.0 mm of depth. Units were sorted and analysed using custom written codes in MATLAB as follows. For every channel, 1.5 ms event waveforms were detected at 1 ms spaced peaks which exceeded 5 times root-mean-square (RMS). The first 3 principle components of the data were then calculated and all waveforms were automatically grouped into 5 clusters using agglomerative hierarchical clustering. These average waveforms of each cluster were then plotted as 3 principle components to manually exclude artefacts. Mean spike rates across trials were smoothed by a 16.7 ms averaging window. The width of the average spike waveforms of each unit was used to exclude narrow spiking neurons (20% narrowest spikes of the sorted data was excluded). To build response profiles, pre-puff spike rate of each unit within 900 ms preceding the puff was subtracted. To find units with an inhibition of the ongoing activity, the response profiles were sorted based on the changes in firing rate (delta firing rate) during stimulus presentation (2 s). We analysed the first and last quartiles (25% top and 25% bottom) of the sorted data in each group which were considered as the “excited cells” and the “inhibited cells” respectively. In order to include the head and whisker motions in the analysis, MATLAB’s foreground detection algorithm was used to obtain a motion index (average of the image in binary foreground images). Spike triggered averaging (STA) of the motion index was then performed for each unit and the resulting STA was normalised within the interval (-3 to 3 seconds). A bootstrapping procedure was performed on the difference of the time-varying signals (firing rat and STA) between cKO and the control. A 95% confidence bound was then calculated at each time point and plotted together with the actual difference between the two groups to show their statistical significance of the differences at each time point. To compare overall spontaneous or evoked spike rates between the two groups, an empirical cumulative distribution was plotted for each condition and each quartile (0-1 s of the first and 0.2-1.2 s of the last quartile to exclude rebound activity and the delay of the suppression). A statistical test was first applied on the overall distributions. Enhanced deviations of the *Er81* cKO firing rate distributions from the control distributions appeared as an intersection of the two cumulative distributions. We found such intersections in the estimated probability density functions of each distribution and used it as a criterion to separate data into two groups for independent statistical analysis. Correlations between firing rate and the air puff and motion were calculated using cross-correlation analysis in MATLAB. For a smooth estimation of the firing rate spike times were convolved by a leaky integrator function (*dy/dt* = −*y*). Onsets of the air puffs were also convolved by the same function. The peaks of cross-correlations were chosen within a ±2 s lag. Since there was no genotype effect on the lag of the peak we used the peak cross-correlation to analyse the strength of correlations between the variables.

### Immunohistochemistry

Animals were perfused transcardially with 0.01M phosphate buffered saline (PBS) to eliminate blood and extraneous material followed by 4% paraformaldehyde (PFA) under isoflurane anaesthesia. Brains were left incubated for 2-5 hours in PFA. The fixative was then removed from tissues by 3 washes in PBS. Tissues were sectioned at 60µm using a Leica 1000S vibratome and kept in a cryoprotective ethylene glycol solution at −20°C until processed for immunofluorescence.

Sections were first washed and permeabilised, then non-specific binding sites were blocked by immersing the tissue in 10% normal donkey serum, 2% BSA in PBS-T for 2 hours. Tissues were then stained using the following primary antibodies overnight: mouse anti-β Galactosidase (1:1000; Promega), rabbit anti-Er81 (1:5000; generous gift from Prof Silvia Arber; KO-validated (Dehorter et al., 2015)), Chicken anti-GFP (1:3000; Aves Lab), mouse anti-parvalbumin (1:3000; Sigma), goat anti-ChAT (1:200; Merck), guinea pig anti-vGluT1 and -vGluT2 (1:2000; Chemicon) and anti-vGluT3 (1:2000; Merck), mouse anti-Lhx6 (1:X; Santa Cruz), rabbit anti-mgluR5 (1:50; Alomone), sheep anti-Neuropeptide Y (1:1000; Merck), rabbit anti-KCNQ2 and anti-HCN2 (1:200; Thermofisher), mouse anti-c-Fos (1:500; Santa Cruz) or rabbit anti-p-Ser240–244-S6rp (1:500; Cell Signalling). After 3 times washes with 15 min intervals, we added anti-rabbit, anti-Chicken, anti-mouse, anti-goat Alexa 488, 555 or 647 (1:200; Life Technologies) secondary antibodies for 2-3 hours or anti-guinea pig biotinylated (1:200; Jackson), anti-sheep biotinylated (1:200; Life technology) followed by Streptavidin 555 or 647 for 1. After 3 times washes with 15 min intervals, slices were stain for 10 min with DAPI (5µM; Sigma), mounted on Livingstone slides then covered with Mowiol (Sigma) and coverslip (Thermofisher).

After patch clamp recordings, slices were immediately fixed in 4% PFA for 2-5 hours, rinsed in PBS (3 times, 30 min intervals) and kept overnight at 4 °C. The same procedure as described above, was performed for immunostaining. Images were acquired by a Nikon A1 confocal laser scanning microscope. Stained sections of control and mutant mice were imaged during the same imaging session and laser power, photomultiplier gain, pinhole and detection filter settings were kept constant. Immunofluorescence signals were quantified using ImageJ or MATLAB using routine particle analysis procedures to obtain soma or nuclear masks. The signal within masks were normalised to the background. The background masks were obtained by segmentation of thick fibres passing the striatum. To quantify the percent of βGal positive cells, the Cartesian distances between the weighted centroids of the ChAT^+^ nuclei masks and βGal masks (threshold at mean + 0.4 standard deviation) were calculated. If the nucleus mask was within 5 µm of a βGal mask, the cell was considered as ChAT^+^ βGal^+^.

### Western Blot

Brains were dissected in ACSF and the striatum and cortex were extracted. Tissues were placed in lysis buffer with 1% ß-mercaptoethanol (Sigma) prior to storage at -80°C. Total protein was extracted using the AllPrep DNA/RNA/Protein Mini Kit (Qiagen #80004) according to manufacturer’s instructions, including DNase treatment. The pellets were suspended in ALO buffer then stored at −20°C. We extracted the striatum and cortex and homogenised by handheld homogeniser, at 1:5 g/mL (50mM Tris–HCl pH 7.4, 0.32M sucrose, 5mM EDTA, 1% Triton X-100, 1% protease inhibitor cocktail) then stored at −80°C. 20µL of each sample was run on 8-16% Mini-PROTEAN® TGX Stain-Free™ Gels (200V, 30 min), transferred to a PVDF membrane (340mA, 70 min). The membrane was then blocked (1H, 3% BSA in TBS-T), incubated overnight with primary antibody overnight with the guinea-pig anti-vGluT3 (Merck; 1:2000), Goat anti-ChAT (Merck; 1:500), Mouse anti-Lhx6 (Santa Cruz Biotechnology; 1:500), then washed (3 times washes with 5 min intervals, TBS-T) before incubation with HRP-conjugated secondary antibody for 2 hours (Biorad, 1:1000). After 3 times washes with 5 min intervals with TBS-T, we developed with Clarity™ Western ECL Blotting Substrates (Biorad). Membranes were imaged by ChemiDoc™ MP System (Biorad). For re-probing, blots were stripped with harsh stripping buffer for 30mins, then re-blocked and probed B-tubulin for 1h with the mouse anti-ß-tubulin (1:1000; Sigma). Bands were analysed by area under curve relative to tubulin loading controls (Image Lab™ software).

### Fluorescence Activated Cell Sorting (FACS)

Mice were rapidly decapitated and brain slices were prepared as for electrophysiological recordings. Dorsal striatum was dissected and placed in ACSF, in a 15ml tube with 10 ml digestion buffer (0.5 g Trehalose and 10 mg Pronase, 1 mg DNAse I, 50 µl MgCl2 1 M in oxygenated ACSF; Sigma) at 37°C for 30 min with inversion every 10 min. Tissues were allowed to settle on ice for 1 min, digestion buffer was removed and the pellet was washed with 2 ml washing buffer (10ml ACSF, 0.5 g Trehalose, 1mg DNAse I, 50 µl MgCl2 1M in oxygenated ACSF) at 4 °C. To dissociate the cells, we resuspended the tissue in 1ml wash buffer and triturated 12-15 times with 1 ml pipette. Tubes were placed on ice, large tissue pieces were allowed to settle, and ∼500 µl of cloudy suspension containing dissociated cells was transferred to a 1.5 ml tube on ice. This process was repeated for the remaining 500 µl suspension with 200 µl pipette 12-15 times and pooled into the 1.5 ml tubes on ice. The last cells were removed by adding 200 µl to the original tube. The cell suspension was filtered through a 70 µm tick mesh filter into a snap cap tube and 200 µl DAPI 5 µM was added to exclude dead cells. FACS Melody software was used for cell sorting. Cells were collected in 350 µl of lysis buffer with 1% B-mercaptoethanol prior to storage at -80 °C for RNA extraction.

### Q-rt PCR

Total RNA was extracted using the RNeasy Micro kit (Qiagen) according to manufacturer’s instructions, including DNase treatment. RNA concentration and purity were determined using the Nanodrop spectrophotometer (Thermo Fisher Scientific). RNA was reverse transcribed into cDNA using SuperScriptIII First Strand Synthesis System (Invitrogen) with random hexanucleotides according to manufacturer instructions. cDNA was analysed using real time qPCR, in technical triplicates using TaqMan™Gene Expression Assay. Reactions were performed on a MicroAmp™ Optical 384-Well Reaction Plate with Barcode (Applied Biosystems), Each 10 µl reaction included 5.5 µl of TaqMan™ Universal PCR Master Mix (Applied Biosystems), 0.5 µl of Taqman probes gene expression assay and 4.5 µl of cDNA (5-10 ng total). PCRs were monitored using the 7900HT Realtime PCR system (Applied Biosystems). The program began with 2 min at 50 °C, 10 min at 95 °C, 40 cycles of 15 s at 95°C and 1 min at 60°C.

### Morphology and synaptic bouton analysis

*Er81^+/+^; Nkx2.1-CreER; RCE-GFP* and *Er81^F/F^; Nkx2.1-CreER; RCE-GFP* P0 mouse pups were injected with low-titer Tamoxifen (Sigma Aldrich, 25 µl of 10 mg/ml solution in corn oil, i.p.) to induce GFP expression and *Er81* conditional knockout in MGE derived interneurons Nkx2.1. Brains were removed after deep anaesthesia under 3-4% isoflurane and PBS followed by 4% PFA was transcardially perfused. We then proceeded to the immunohistochemistry to select Nkx2.1-GFP/ChAT^+^ interneurons. Images were acquired using a Nikon A1 Confocal microscope and the Nikon Instruments Elements software. Image stacks were acquired for morphological reconstruction with 20x objective. Images of neurons were acquired using with 60x objective and 2x scanning zoom for bouton analysis (stacks of 20-25 µm with 1µm resolution). Parameters were kept consistent across wild type and cKO slices.

Morphological quantification was done using the Surface and Filament tools in the IMARIS software. The automatic component of the filament tool reconstructed the dendritic field of the neuron with the dendrite beginning point set to 15µm and the end point set to 1µm. The volume of dendritic spread was found by using the Convex Hull Xtension of IMARIS. Bouton analysis was done using the Surface and Spots tools in IMARIS. A co-localisation channel was created to identify individual boutons. This channel was then passed by a threshold to filter weak co-localisations. The brightest 30% of co-localised particles were chosen as boutons. Spots outside 1.5 µm of the neuron surface were also excluded. Bouton density was calculated as the number of boutons divided by the surface area of the soma.

### Behavioural task

6 weeks-old *ChAT-Cre; Er81^+/+^* (control) and *ChAT-Cre; Er81^flox/flox^* (Er81 cKO) mice were caged in standardised room conditions under a 12:12 light: dark cycle with ad libitum food and water. Starting 5 days before the behavioural task and throughout the training, mice were maintained under a mild water restriction (2 h ON/ 22 h OFF) with *ad libitum* food access. Mice were then trained in an arena (40 cm length × 27 cm width × 25 cm height) made of transparent plexiglass, divided into three 9 cm corridors, and placed on top of a DigiGait apparatus (Mouse Specifics, Inc.) where a camera recorded the ventral view of the animal in the middle corridor. The outer corridors were equipped with dispensers to deliver a drop of 10% sucrose in correct trials. We used custom-written MATLAB programs to automatically deliver rewards upon lick detection through capacitance sensors, to monitor mouse behaviour and to extract the data.

At the beginning of a session, mice were placed at the starting point of the middle corridor and had to go through one of the one-way gates to choose between the two outer corridors. If the choice was correct, the mouse could collect the reward automatically delivered by the dispenser. The mouse could then go back to the start area through a second one-way gate to start the next trial. Prior to acquisition, mice were habituated to the apparatus over a period of six days during which they were allowed to move freely in the maze, with both dispensers delivering the reward. To complete the habituation session, mice had to complete 10 trials, one trial being defined as running from the start site to one of the outer corridors, then back to the start site. If 10 trials were not completed after 40 minutes the session was terminated.

During the acquisition phase, each mouse was required to complete 20 trials per session and one session a day for 3 days. Only one side contained the reward. The rewarding side (left or right) was homogenised within an experimental group and balanced between groups. On Day 4, corresponding to reversal 1, the side containing the reward was switched for each mouse. Mice were then exposed to the new side for 10 consecutive sessions (Day 4 to Day 13). From Day 14, corresponding to reversal 2, the reward sides were switched again. Mice were tested for 3 days. After the training in Day 4 and Day 14, some mice were removed from training for immunohistochemistry analyses (Day 4: n = 6 control and 4 Er81 mutant mice; Day 14: n = 5 control and 4 Er81 mutant mice). Performance was defined as the percentage choice of the reward side, regardless of reward collection by the animal.

### Statistics

Statistical analysis was carried out in MATLAB or Prism software. Data obtained for the expression of protein within single cells was analysed with a D’Agostino and Pearson normality test to determine if the assumption of a normal distribution was met. If the data passed this test (and was normal) a non-paired, two tailed t-test was applied when comparing two groups. And an ANOVA (Analysis of Variance), and then multiple t-tests when comparing more than two groups. If the data failed this test, and thus did not follow a normal distribution, a Mann-Whitney or Wilcoxon rank sum test was performed to compare two groups. Correlative data was passed through correlation analysis in Prism to generate a p-value based on the Pearson correlation coefficient.

In case multiple tests were done on a specific measurement, a Benjamini-Hochberg procedure was followed to correct for multiple comparison errors. P values < 0.05 were considered statistically significant. Data are presented as mean and SEM. In the case of bar and whisker plots, the data is represented as median and interquartile range (25-75% of the data as the bar and the whole data range as whiskers). Multiple linear regression test (using fitlm function in MATLAB) was applied to test the interactions between two independent variables.

## SUPPLEMENTAL INFORMATION

Supplemental Information includes 5 figures and 3 tables.

## ACKNOWLEDGEMENTS

We would like to thank Prof Arber and Prof Marin for the Er81 antibody and Er81 conditional mutant mice. This work was supported by the National Health and Medical Research Council (grant APP1144145) and the Australian National University.

## AUTHORS CONTRIBUTIONS

N.D., Y.R-S. and E.A. designed the experiments. Y.R-S. performed and analysed the electrophysiological recordings. L.G. & AS. A performed the behavioural experiment. L.G., AS. A, and Y.R-S analysed the behavioural data. Y.R-S., NY.A., A.R.C., AS. A performed and analysed immunohistochemistry data. Y.S. and A.R.C. performed the experiments in molecular biology. Y.R-S., AS. A and N.D wrote the manuscript.

## SUPPLEMENTARY FIGURES

**Supplementary figure 1:**
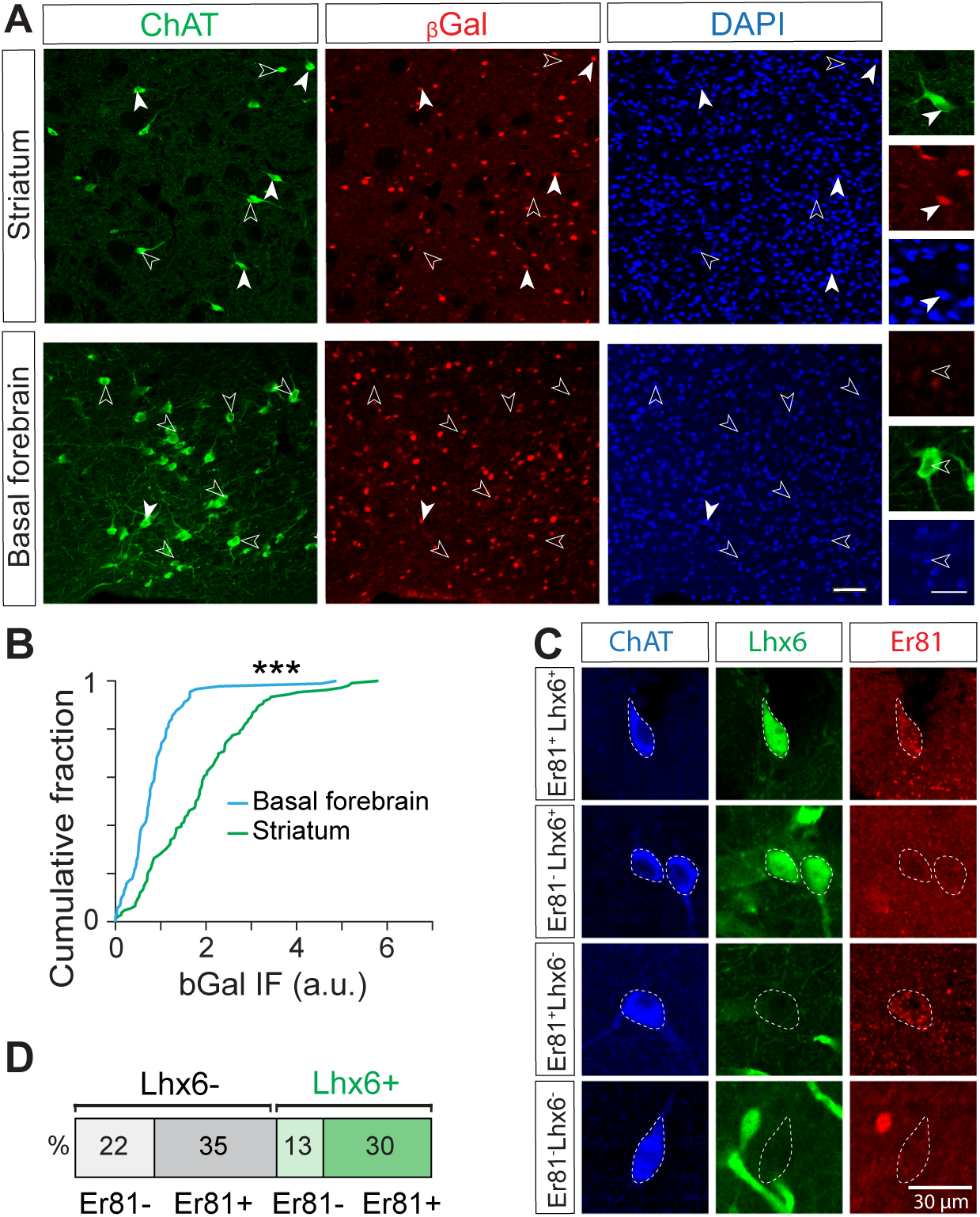
Pattern of expression of the *Er81* and *Lhx6 transcription factors* in cholinergic neurons. **A.** βGal expression in ChAT^+^ neurons in the striatum (top) and basal forebrain (bottom). Scale bars: 50 µm and 25 µm (inset). **B.** Cumulative distribution of the βGal-expressing ChAT^+^ cells in the striatum (green, n = 106 cells) and basal forebrain (blue, n = 89 cells) (**\*\*\*** p < 0.001). **C.** Immunostaining for ChAT (blue), *Lhx6* (GFP, green) and Er81 (red). **D.** Classification of CINs into 4 groups based on Er81 and *Lhx6* expressions.

**Supplementary figure 2:**
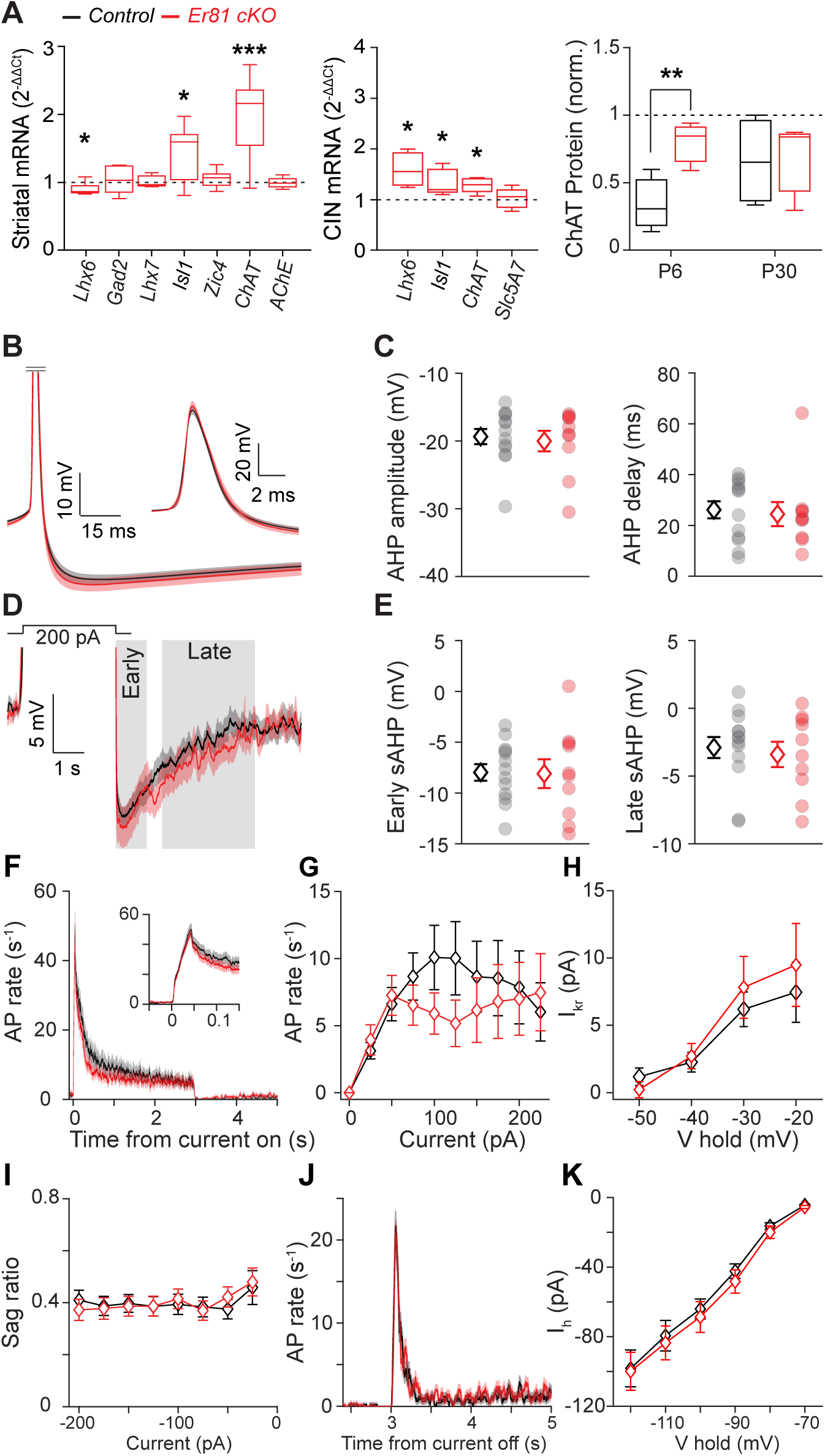
Properties of the P6 cholinergic interneurons in the absence of *Er81*. **A.** mRNA levels in P6 striatum (left) and CINs (middle) in the absence of *Er81*, relative to the control (dashed line; n = 5-12 mice). Expression of ChAT in CINs (right, P6; control; n = 5, *Er81* cKO n = 6, P30; control; n = 4 per group, p = 0.730) **B.** Average spike waveforms (± SEM) in the CINs of the control (black, n = 13, 4 mice) and the *Er81* cKO (red, n = 10, 3 mice) at P6. **C.** AHP properties in control and the *Er81* cKO mice (Left: AHP amplitude, p = 0.780. Right: AHP delay, p = 0.476, sample size as in B). **D.** Average membrane potential across CINs after AP truncation. Grey shadings: Early and late phases of slow AHP (sAHP, sample size as in B). **E.** Average of the early (Left; p = 0.935) and late phase of sAHP (right; p = 0.674) in the two conditions (sample size as in B). **F.** Average firing rate across CINs and all positive current steps in the control (black) and the *Er81* cKO CINs (red, 0.1-3 s; p = 0.142). Inset is an expanded view on the time axis (0-0.1 s; p = 0.130). **G.** Firing rate during positive current steps across control (n = 16, 5 mice) and the *Er81* cKO (n = 17 cells, 4 mice) CINs at P6 (p ≥ 0.121 in all conditions). **H.** Average I_Kr_ amplitude recorded in voltage clamp mode (n = 9 control, 5 mice and 10 cKO 3 mice CINs, p ≥ 0.400 in all conditions). **I.** Voltage sag ratio at negative current steps (n = 16 control, 5 mice and n = 17 cKO CINs; p ≥ 0.242 in all conditions). **J.** Firing rate at the offset of negative current steps (3-3.23 s; p = 0.957, 3.23-5 s; p = 0.427). **K.** The average I_h_ recorded in voltage clamp mode (from −120 mV to −70 mV) in control (n = 13 cells, 5 mice) and cKO (n = 11 cells, 4 mice, p ≥ 0.305 in all conditions) CINs. Data are presented as mean ± SEM (diamond). Statistical tests are t-test or Wilcoxon rank-sum depending on data normality. Multiple comparisons are corrected using Benjamini-Hochberg procedure. **\*** p < 0.05, **\*\*** p < 0.01, **\*\*\*** p < 0.001.

**Supplementary figure 3:**
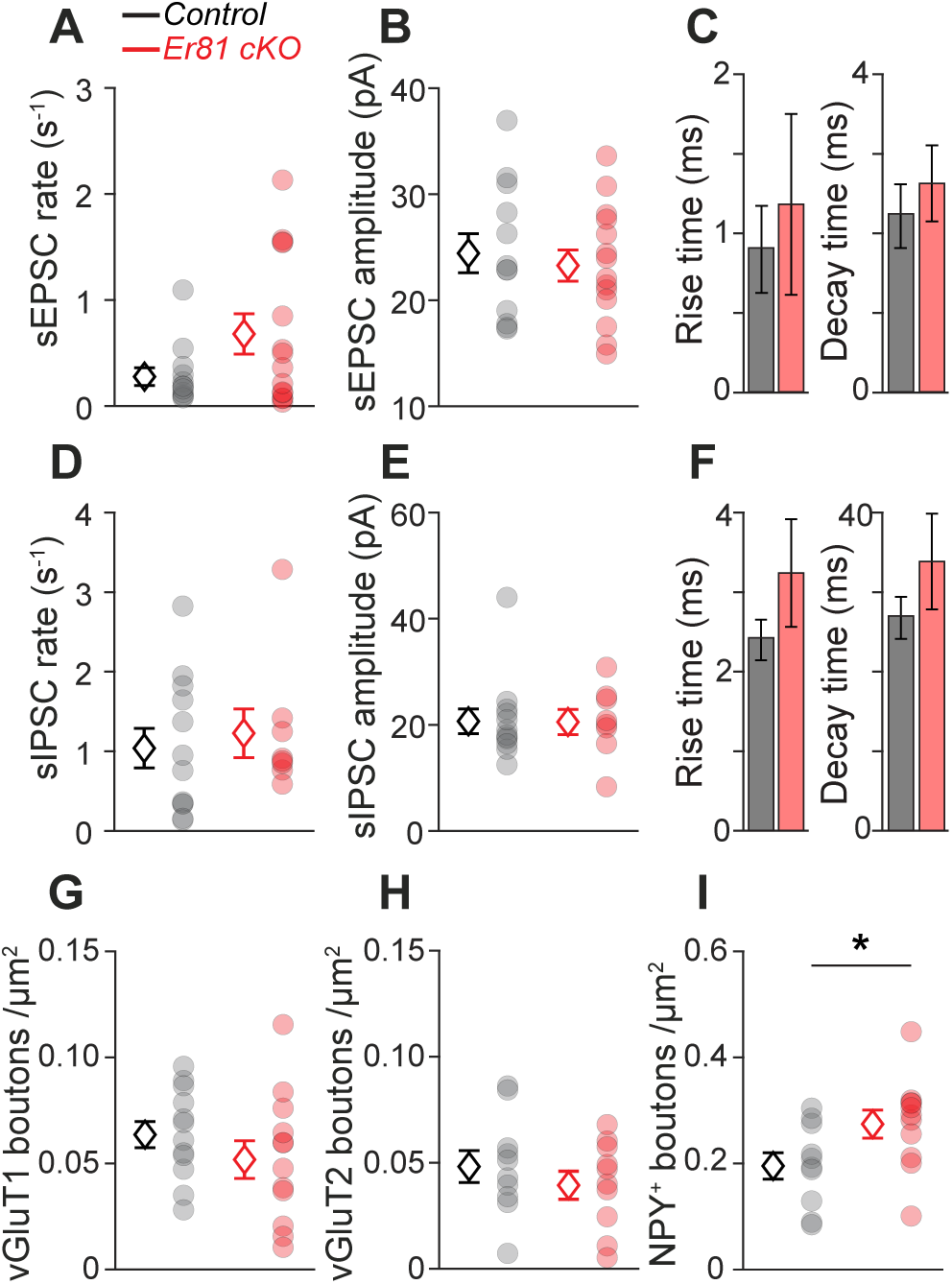
Analysis of the synaptic inputs of P6 CINs revealed no functional change in the absence of *Er81*. **A-C.** Rate (**A**, p = 0.368), amplitude (**B**, p = 0.624) rise time (**C**, left, p = 0.777) and decay time (**C,** right, p = 0.456) of spontaneous excitatory postsynaptic currents (sEPSCs) across CINs (control; n = 12, 6 mice, *Er81* cKO; n = 14, 3 mice). **D-F.** Rate (**D**, p = 0.563), amplitude (**E**, p = 0.512) rise time (**F**, left, p = 0.199) and decay time (**F,** right, p = 0.263) of spontaneous inhibitory postsynaptic currents (sIPSCs) across CINs (control; n = 12 cells, *Er81* cKO; n = 8 cells). **G.** Quantification of vGluT1 bouton density (control, n = 12 cells, 3 mice, *Er81* cKO, n = 12 cells, 3 mice, p = 0.288). **H.** Density of vGluT2 boutons (control, n = 10 cells, 2 mice, *Er81* cKO, n = 10 cells, 3 mice, p = 0.392). **I.** NPY^+^ bouton density (control, n = 10 cells, 2 mice, *Er81* cKO, n = 11 cells, 3 mice). Data are presented as means ± SEM. **\*** p < 0.05. Averages are shown as diamonds and individual neurons are plotted as grey (control) and red (*Er81* cKO) circles.

**Supplementary figure 4:**
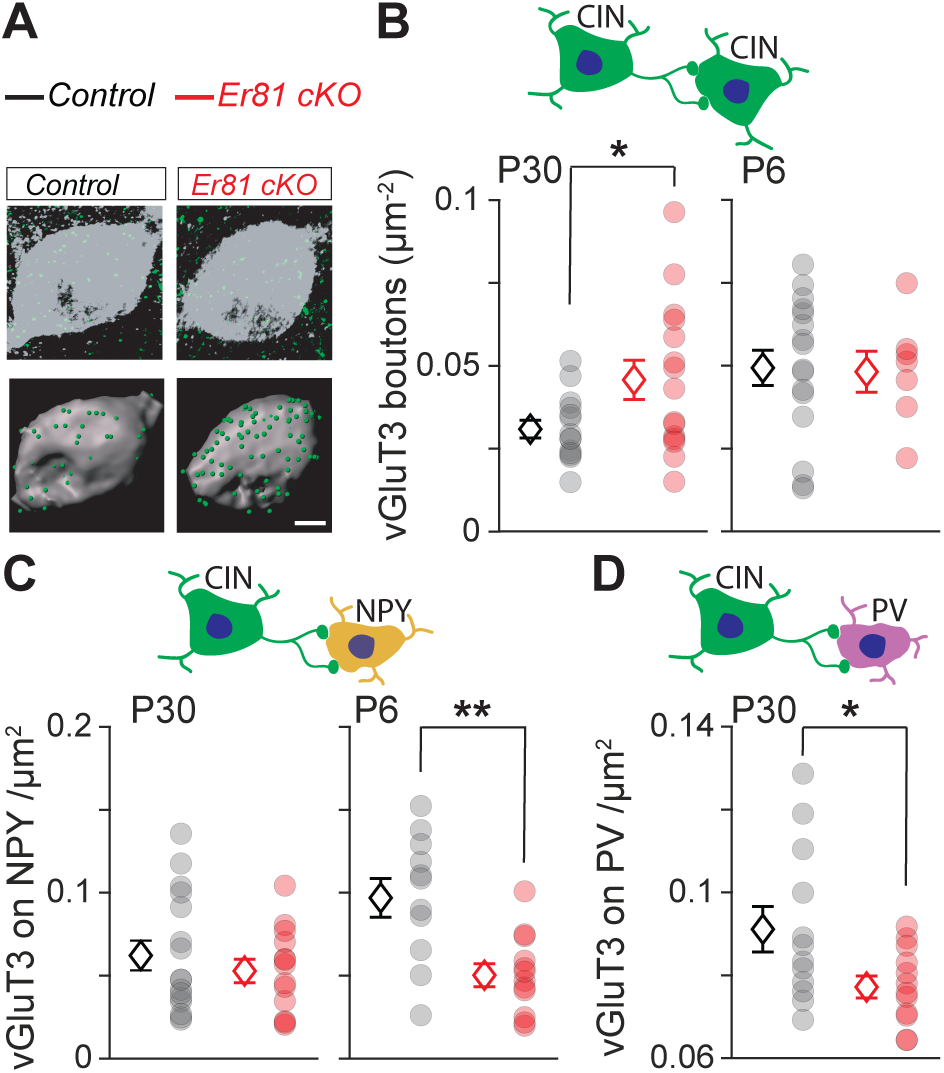
Cholinergic synapses are reduced in the absence of *Er81*. **A.** Example CINs (grey) stained for vesicular glutamate transporter 3 (vGluT3, green) representing cholinergic boutons (top; raw image, bottom; 3D reconstruction). **B.** Quantification of vGluT3 bouton density on CINs at P30 (left; control, n = 14 cells, 3 mice; *Er81* cKO, n = 15 cells, 3 mice) and P6 (right, control; n= 16 cells, 3 mice, *Er81* cKO; n = 7 cells, 3 mice, p = 0.899). **C.** Quantification of vGluT3 bouton density on the NPY^+^ interneurons at P30 (left, control; n = 16 cells, 2 mice, *Er81* cKO; n = 13 cells, 2 mice, p = 0.438) and P6 (right; control, n = 11 cells, 2 mice, cKO, n = 12 cells, 3 mice). **D.** Quantification of vGluT3 bouton density onto PV^+^ interneurons at P30 (control; n = 12 cells, 3 mice, *Er81* cKO; n = 14 cells, 3 mice). Data are presented as means ± SEM. **\*** p < 0.05, **\*\*** p < 0.01.

**Supplementary figure 5:**
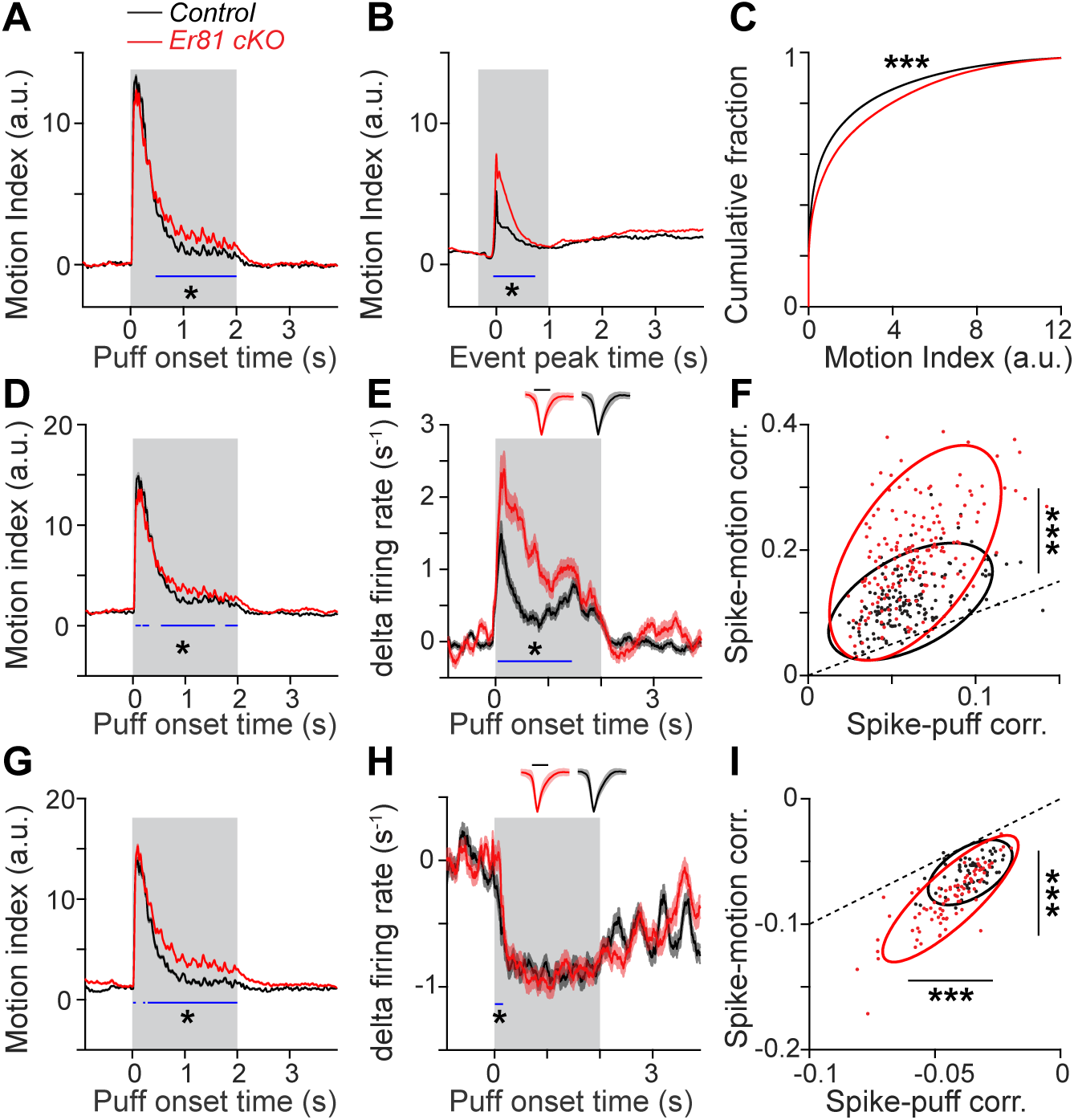
Enhanced correlations between striatal firing and sensorimotor inputs in the *Er81* cKO mice. **A.** Whisker and nose movements (motion index) relative to the puff train onset for all units from the control (n = 731) and the *Er81* cKO mice (n = 732). **B.** Voluntary movements outside the air puff period for all units, aligned to the peaks of motions detected at 0.7 times RMS of the whole trace of the motion index. **C.** Cumulative distribution of the motion indices throughout the recordings (control; n = 21, 3 mice, *Er81* cKO; n = 23, 4 mice). **D.** Motions relative to the puff train onset for the excited units (corresponding to units in Figure 6 D) **E.** Average delta firing rate across the excited units (units in Figure 6D). The inset shows the normalised spike waveforms (scale bar; 500 ms). **F.** Scatter plots of the peak cross-correlation between the firing rate of excited units and their corresponding variables of air puff (x-axis) or motion (y-axis). Dots represent individual units with positive correlation regarding both variables and the ellipses are the first contours of the fitted Gaussian mixture models (control; n = 161, 3 mice, *Er81* cKO; n = 173, 4 mice). **G.** Grand average of the motions relative to the puff train onset for the inhibited units (corresponding to Figure 6 F) **H.** Average delta firing rate of the inhibited units (corresponding to Figure 6 F). The inset shows the average spike waveforms (amplitude is normalised) of the inhibited units of the two groups (scale bar; 500 ms). **I.** Peak cross-correlations between the firing rate of the inhibited units and the stimulus (x-axis) or the motion (y-axis). Dots represent individual units with a negative correlation regarding both variables and the ellipses are the first contours of the fitted Gaussian mixture models (control; n = 66, 3 mice, *Er81* cKO; n = 88, 4 mice). Traces are presented as means ± SEM. Blue lines show significant time points in the shaded regions (two-sided permutation test). **\*** p < 0.05, **\*\*\*** p < 0.001.

**Table S1:**
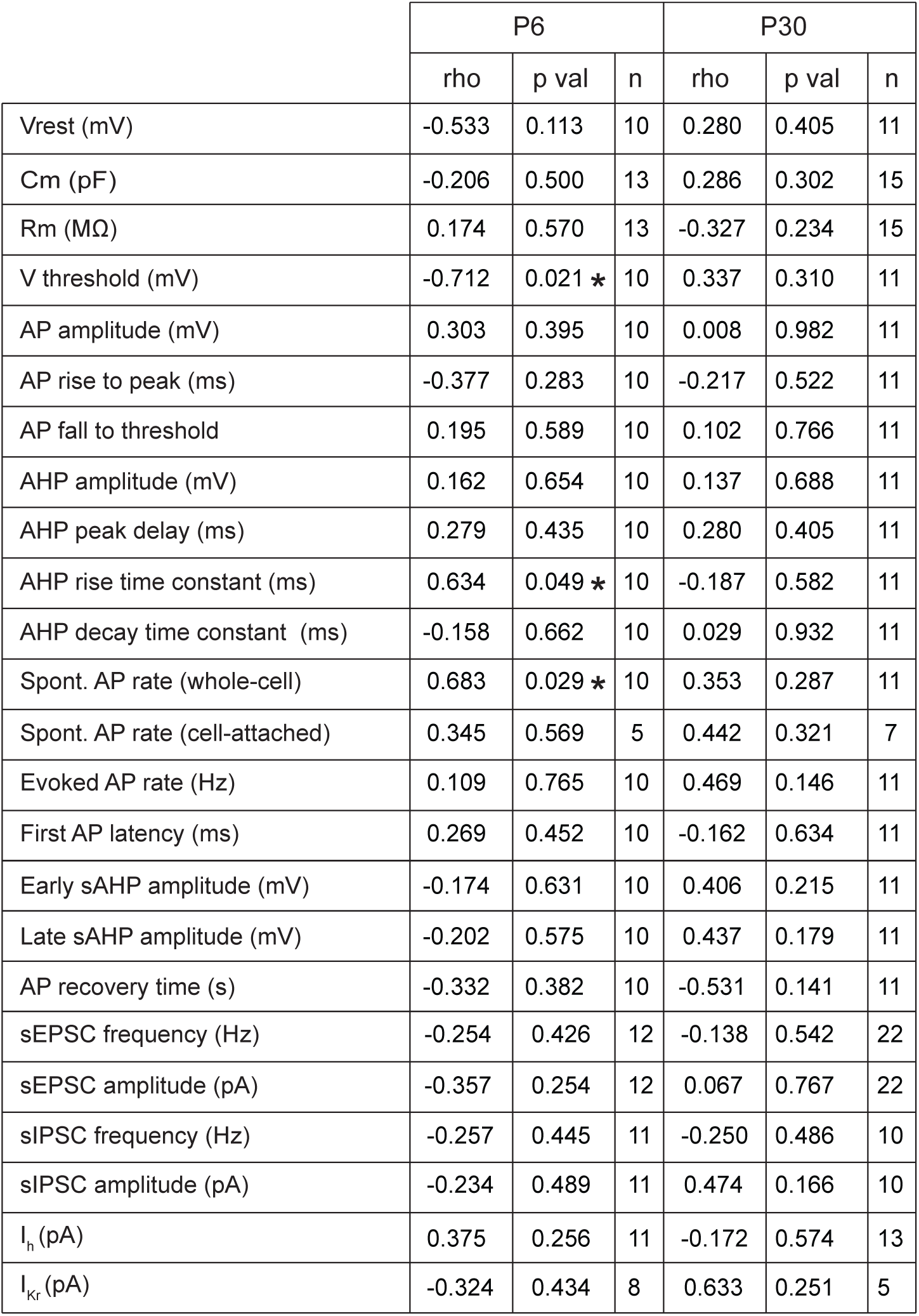
Lack of correlations between most of the physiological parameters and the Er81 expression levels in striatal CINs at P6 and P30. Vrest; resting membrane potential, Cm; membrane capacitance, Rm; membrane resistance, V threshold; threshold potential, AP; action potential, AHP; afterhyperpolarisation, Spont.; spontaneous, sAPH; slow AHP. sEPSC; spontaneous excitatory postsynaptic currents, sIPSC; spontaneous inhibitory postsynaptic currents. I_h_; hyperpolarization-activated current, I_Kr_; delayed rectifier current. **\*** p < 0.05, Pearson’s correlation.

**Table S2:**
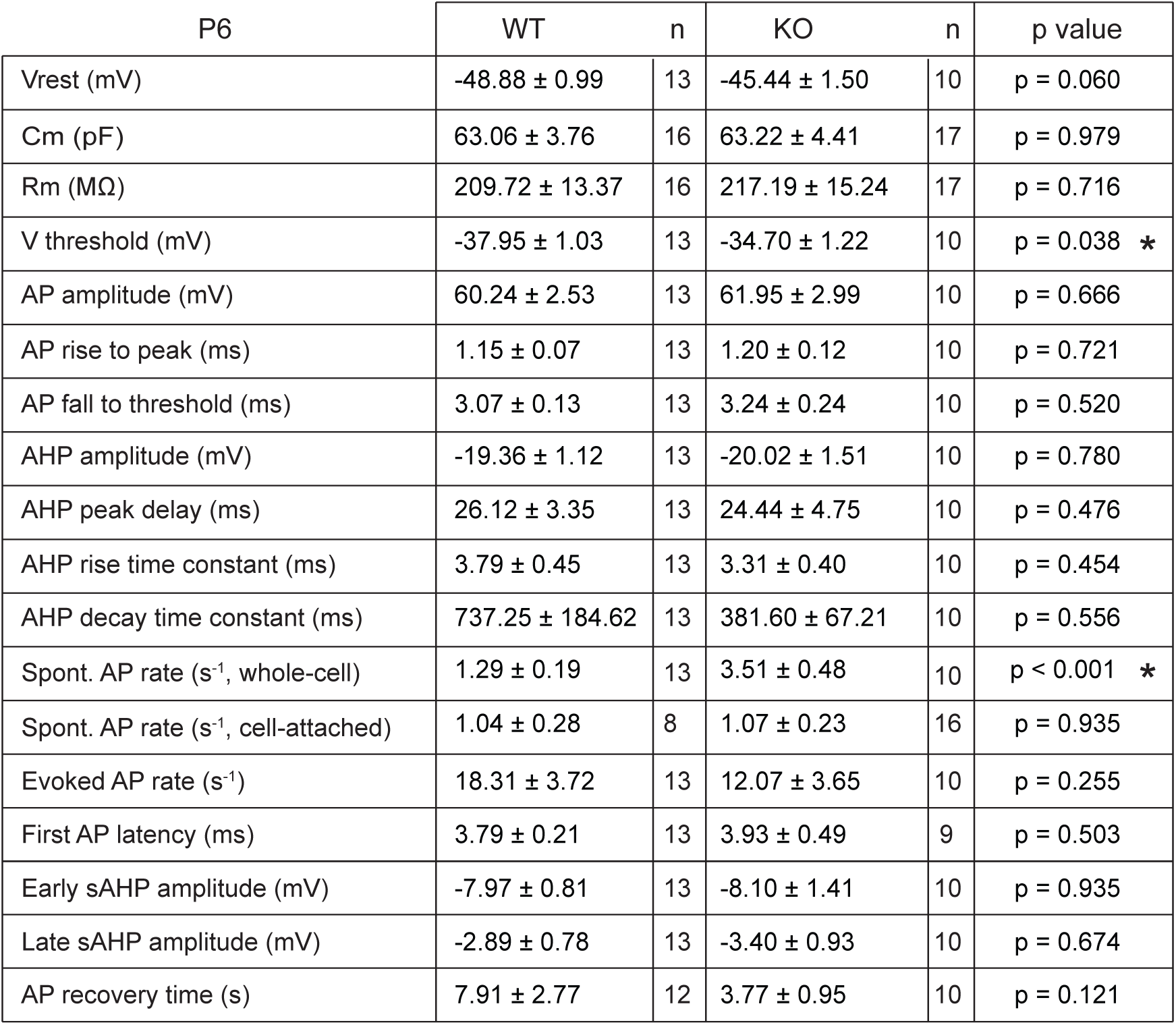
Intrinsic properties of striatal CINs in the *Er81* cKO and the control mice at P6. Vrest; resting membrane potential, Cm; membrane capacitance, Rm; membrane resistance, V threshold; threshold potential, AP; action potential, AHP; afterhyperpolarisation, Spont.; spontaneous, sAPH; slow AHP. Data are presented as mean ± SEM. **\*** p < 0.05.

**Table S3:**
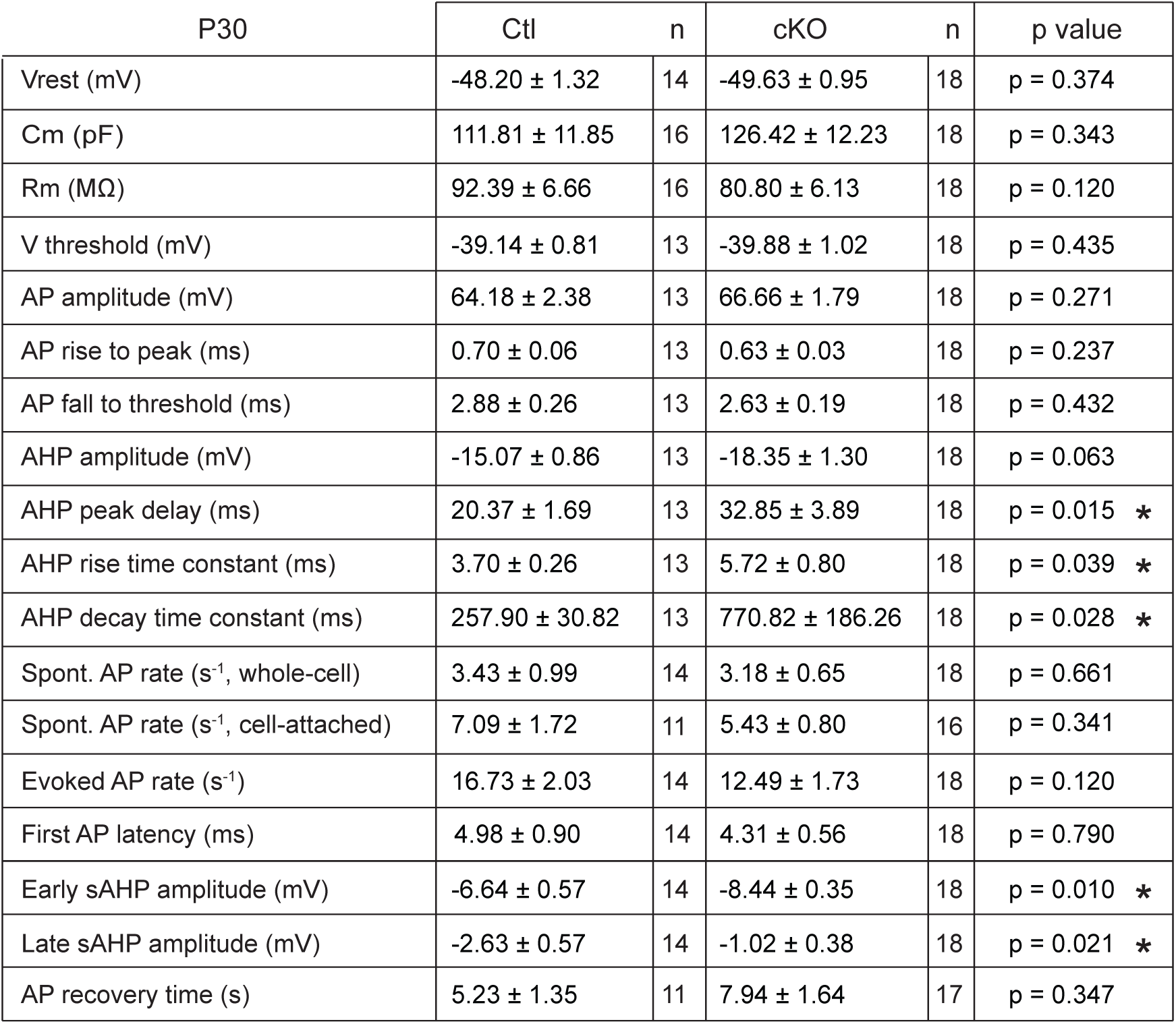
Intrinsic properties of striatal CINs in the *Er81* cKO and the control mice at P30. Vrest; resting membrane potential, Cm; membrane capacitance, Rm; membrane resistance, V threshold; threshold potential, AP; action potential, AHP; afterhyperpolarisation, Spont.; spontaneous, sAPH; slow AHP. Data are presented as mean ± SEM. **\*** p < 0.05.

## REFERENCES

1. Abe, H., Okazawa, M. & Nakanishi, S. 2011. The Etv1/Er81 transcription factor orchestrates activity-dependent gene regulation in the terminal maturation program of cerebellar granule cells. Proc Natl Acad Sci U S A, 108, 12497–502.

2. Ahmed, N. Y., Knowles, R. & Dehorter, N. 2019. New Insights Into Cholinergic Neuron Diversity. Front Mol Neurosci, 12, 204.

3. Allaway, K. C. & Machold, R. 2017. Developmental specification of forebrain cholinergic neurons. Dev Biol, 421, 1–7.

4. Alvarez-Baron, C. P., Klenchin, V. A. & Chanda, B. 2018. Minimal molecular determinants of isoform-specific differences in efficacy in the Hcn channel family. J Gen Physiol, 150, 1203–1213.

5. Aoki, S., Liu, A. W., Akamine, Y., Zucca, A., Zucca, S. & Wickens, J. R. 2018. Cholinergic interneurons in the rat striatum modulate substitution of habits. Eur J Neurosci, 47, 1194–1205.

6. Aoki, S., Liu, A. W., Zucca, A., Zucca, S. & Wickens, J. R. 2015. Role of Striatal Cholinergic Interneurons in Set-Shifting in the Rat. J Neurosci, 35, 9424–31.

7. Apicella, P. 2017. The role of the intrinsic cholinergic system of the striatum: What have we learned from Tan recordings in behaving animals? Neuroscience, 360, 81–94.

8. Arber, S., Ladle, D. R., Lin, J. H., Frank, E. & Jessell, T. M. 2000. Ets gene Er81 controls the formation of functional connections between group Ia sensory afferents and motor neurons. Cell, 101, 485–98.

9. Balleine, B. W. 2019. The Meaning of Behavior: Discriminating Reflex and Volition in the Brain. Neuron, 104, 47–62.

10. Bennett, B. D., Callaway, J. C. & Wilson, C. J. 2000. Intrinsic membrane properties underlying spontaneous tonic firing in neostriatal cholinergic interneurons. J Neurosci, 20, 8493–503.

11. Brimblecombe, K. R., Threlfell, S., Dautan, D., Kosillo, P., Mena-Segovia, J. & Cragg, S. J. 2018. Targeted Activation of Cholinergic Interneurons Accounts for the Modulation of Dopamine by Striatal Nicotinic Receptors. eNeuro, 5.

12. Cave, J. W., Akiba, Y., Banerjee, K., Bhosle, S., Berlin, R. & Baker, H. 2010. Differential regulation of dopaminergic gene expression by Er81. J Neurosci, 30, 4717–24.

13. Cui, Y., Prokin, I., Mendes, A., Berry, H. & Venance, L. 2018. Robustness of Stdp to spike timing jitter. Sci Rep, 8, 8139.

14. Dehorter, N., Ciceri, G., Bartolini, G., Lim, L., Del Pino, I. & Marin, O. 2015. Tuning of fast-spiking interneuron properties by an activity-dependent transcriptional switch. Science, 349, 1216–20.

15. Ding, B., Cave, J. W., Dobner, P. R., Mullikin-Kilpatrick, D., Bartzokis, M., Zhu, H., Chow, C. W., Gronostajski, R. M. & Kilpatrick, D. L. 2016. Reciprocal autoregulation by Nfi occupancy and Etv1 promotes the developmental expression of dendrite-synapse genes in cerebellar granule neurons. Mol Biol Cell, 27, 1488–99.

16. Ding, J. B., Guzman, J. N., Peterson, J. D., Goldberg, J. A. & Surmeier, D. J. 2010. Thalamic gating of corticostriatal signaling by cholinergic interneurons. Neuron, 67, 294–307.

17. Doig, N. M., Magill, P. J., Apicella, P., Bolam, J. P. & Sharott, A. 2014. Cortical and thalamic excitation mediate the multiphasic responses of striatal cholinergic interneurons to motivationally salient stimuli. J Neurosci, 34, 3101–17.

18. Doitsidou, M., Flames, N., Topalidou, I., Abe, N., Felton, T., Remesal, L., Popovitchenko, T., Mann, R., Chalfie, M. & Hobert, O. 2013. A combinatorial regulatory signature controls terminal differentiation of the dopaminergic nervous system in C. elegans. Genes Dev, 27, 1391–405.

19. English, D. F., Ibanez-Sandoval, O., Stark, E., Tecuapetla, F., Buzsaki, G., Deisseroth, K., Tepper, J. M. & Koos, T. 2011. Gabaergic circuits mediate the reinforcement-related signals of striatal cholinergic interneurons. Nat Neurosci, 15, 123–30.

20. Falkenburger, B. H., Jensen, J. B. & Hille, B. 2010. Kinetics of Pip2 metabolism and Kcnq2/3 channel regulation studied with a voltage-sensitive phosphatase in living cells. J Gen Physiol, 135, 99–114.

21. Fino, E., Deniau, J. M. & Venance, L. 2008. Cell-specific spike-timing-dependent plasticity in Gabaergic and cholinergic interneurons in corticostriatal rat brain slices. J Physiol, 586, 265–82.

22. Flames, N. & Hobert, O. 2009. Gene regulatory logic of dopamine neuron differentiation. Nature, 458, 885–9.

23. Flames, N., Pla, R., Gelman, D. M., Rubenstein, J. L., Puelles, L. & Marin, O. 2007. Delineation of multiple subpallial progenitor domains by the combinatorial expression of transcriptional codes. J Neurosci, 27, 9682–95.

24. Flandin, P., Zhao, Y., Vogt, D., Jeong, J., Long, J., Potter, G., Westphal, H. & Rubenstein, J. L. 2011. Lhx6 and Lhx8 coordinately induce neuronal expression of Shh that controls the generation of interneuron progenitors. Neuron, 70, 939–50.

25. Fragkouli, A., Van Wijk, N. V., Lopes, R., Kessaris, N. & Pachnis, V. 2009. Lim homeodomain transcription factor-dependent specification of bipotential Mge progenitors into cholinergic and Gabaergic striatal interneurons. Development, 136, 3841–51.

26. Franklin, N. T. & Frank, M. J. 2015. A cholinergic feedback circuit to regulate striatal population uncertainty and optimize reinforcement learning. Elife, 4.

27. Granger, A. J., Mulder, N., Saunders, A. & Sabatini, B. L. 2016. Cotransmission of acetylcholine and Gaba. Neuropharmacology, 100, 40–6.

28. Graybiel, A. M., Aosaki, T., Flaherty, A. W. & Kimura, M. 1994. The basal ganglia and adaptive motor control. Science, 265, 1826–31.

29. Graybiel, A. M. & Grafton, S. T. 2015. The striatum: where skills and habits meet. Cold Spring Harb Perspect Biol, 7, a021691.

30. Gritton, H. J., Howe, W. M., Romano, M. F., Difeliceantonio, A. G., Kramer, M. A., Saligrama, V., Bucklin, M. E., Zemel, D. & Han, X. 2019. Unique contributions of parvalbumin and cholinergic interneurons in organizing striatal networks during movement. Nat Neurosci, 22, 586–597.

31. He, M., Tucciarone, J., Lee, S., Nigro, M. J., Kim, Y., Levine, J. M., Kelly, S. M., Krugikov, I., Wu, P., Chen, Y., Gong, L., Hou, Y., Osten, P., Rudy, B. & Huang, Z. J. 2016. Strategies and Tools for Combinatorial Targeting of Gabaergic Neurons in Mouse Cerebral Cortex. Neuron, 92, 555.

32. Higley, M. J., Gittis, A. H., Oldenburg, I. A., Balthasar, N., Seal, R. P., Edwards, R. H., Lowell, B. B., Kreitzer, A. C. & Sabatini, B. L. 2011. Cholinergic interneurons mediate fast VgluT3-dependent glutamatergic transmission in the striatum. PloS One, 6, e19155.

33. Hippenmeyer, S., Vrieseling, E., Sigrist, M., Portmann, T., Laengle, C., Ladle, D. R. & Arber, S. 2005. A developmental switch in the response of Drg neurons to Ets transcription factor signaling. PloS Biol, 3, e159.

34. Inokawa, H., Yamada, H., Matsumoto, N., Muranishi, M. & Kimura, M. 2010. Juxtacellular labeling of tonically active neurons and phasically active neurons in the rat striatum. Neuroscience, 168, 395–404.

35. Karvat, G. & Kimchi, T. 2014. Acetylcholine elevation relieves cognitive rigidity and social deficiency in a mouse model of autism. Neuropsychopharmacology, 39, 831–40.

36. Kim, K. S., Duignan, K. M., Hawryluk, J. M., Soh, H. & Tzingounis, A. V. 2016. The Voltage Activation of Cortical Kcnq Channels Depends on Global Pip2 Levels. Biophys J, 110, 1089–98.

37. Klug, J. R., Engelhardt, M. D., Cadman, C. N., Li, H., Smith, J. B., Ayala, S., Williams, E. W., Hoffman, H. & Jin, X. 2018. Differential inputs to striatal cholinergic and parvalbumin interneurons imply functional distinctions. Elife, 7.

38. Kosillo, P., Zhang, Y. F., Threlfell, S. & Cragg, S. J. 2016. Cortical Control of Striatal Dopamine Transmission via Striatal Cholinergic Interneurons. *Cereb Cortex*. Lee, K., Holley, S. M., Shobe, J. L., Chong, N. C., Cepeda, C., Levine, M. S. &

39. Masmanidis, S. C. 2017. Parvalbumin Interneurons Modulate Striatal Output and Enhance Performance during Associative Learning. Neuron, 93, 1451–1463 e4.

40. Lewis, M. & Kim, S. J. 2009. The pathophysiology of restricted repetitive behavior. J Neurodev Disord, 1, 114–32.

41. Lim, S. A., Kang, U. J. & Mcgehee, D. S. 2014. Striatal cholinergic interneuron regulation and circuit effects. Front Synaptic Neurosci, 6, 22.

42. Liodis, P., Denaxa, M., Grigoriou, M., Akufo-Addo, C., Yanagawa, Y. & Pachnis, V. 2007. Lhx6 activity is required for the normal migration and specification of cortical interneuron subtypes. J Neurosci, 27, 3078–89.

43. Lloret-Fernandez, C., Maicas, M., Mora-Martinez, C., Artacho, A., Jimeno-Martin, A., Chirivella, L., Weinberg, P. & Flames, N. 2018. A transcription factor collective defines the Hsn serotonergic neuron regulatory landscape. Elife, 7.

44. Lopes, R., Verhey Van Wijk, N., Neves, G. & Pachnis, V. 2012. Transcription factor Lim homeobox 7 (Lhx7) maintains subtype identity of cholinergic interneurons in the mammalian striatum. Proc Natl Acad Sci U S A, 109, 3119–24.

45. Lozovaya, N., Eftekhari, S., Cloarec, R., Gouty-Colomer, L. A., Dufour, A., Riffault, B., Billon-Grand, M., Pons-Bennaceur, A., Oumar, N., Burnashev, N., Ben-Ari, Y. & Hammond, C. 2018. Gabaergic inhibition in dual-transmission cholinergic and Gabaergic striatal interneurons is abolished in Parkinson disease. Nat Commun, 9, 1422.

46. Magno, L., Barry, C., Schmidt-Hieber, C., Theodotou, P., Hausser, M. & Kessaris, N. 2017. Nkx2-1 Is Required in the Embryonic Septum for Cholinergic System Development, Learning, and Memory. Cell Rep, 20, 1572–1584.

47. Mallet, N., Le Moine, C., Charpier, S. & Gonon, F. 2005. Feedforward inhibition of projection neurons by fast-spiking Gaba interneurons in the rat striatum in vivo. J Neurosci, 25, 3857–69.

48. Mamaligas, A. A., Barcomb, K. & Ford, C. P. 2019. Cholinergic Transmission at Muscarinic Synapses in the Striatum Is Driven Equally by Cortical and Thalamic Inputs. Cell Rep, 28, 1003–1014 e3.

49. Mamaligas, A. A. & Ford, C. P. 2016. Spontaneous Synaptic Activation of Muscarinic Receptors by Striatal Cholinergic Neuron Firing. Neuron, 91, 574–86.

50. Markowitz, J. E., Gillis, W. F., Beron, C. C., Neufeld, S. Q., Robertson, K., Bhagat, N. D., Peterson, R. E., Peterson, E., Hyun, M., Linderman, S. W., Sabatini, B. L. & Datta, S. R. 2018. The Striatum Organizes 3D Behavior via Moment-to-Moment Action Selection. Cell, 174, 44–58 e17.

51. Martos, Y. V., Braz, B. Y., Beccaria, J. P., Murer, M. G. & Belforte, J. E. 2017. Compulsive Social Behavior Emerges after Selective Ablation of Striatal Cholinergic Interneurons. J Neurosci, 37, 2849–2858.

52. Mi, D., Li, Z., Lim, L., Li, M., Moissidis, M., Yang, Y., Gao, T., Hu, T. X., Pratt, T., Price, D. J., Sestan, N. & Marin, O. 2018. Early emergence of cortical interneuron diversity in the mouse embryo. Science, 360, 81–85.

53. Morris, G., Arkadir, D., Nevet, A., Vaadia, E. & Bergman, H. 2004. Coincident but distinct messages of midbrain dopamine and striatal tonically active neurons. Neuron, 43, 133–43.

54. Munoz-Manchado, A. B., Bengtsson Gonzales, C., Zeisel, A., Munguba, H., Bekkouche, B., Skene, N. G., Lonnerberg, P., Ryge, J., Harris, K. D., Linnarsson, S. & Hjerling-Leffler, J. 2018. Diversity of Interneurons in the Dorsal Striatum Revealed by Single-Cell Rna Sequencing and PatchSeq. Cell Rep, 24, 2179–2190 e7.

55. Nelson, A. B., Bussert, T. G., Kreitzer, A. C. & Seal, R. P. 2014. Striatal cholinergic neurotransmission requires Vglut3. J Neurosci, 34, 8772–7.

56. Niswender, C. M. & Conn, P. J. 2010. Metabotropic glutamate receptors: physiology, pharmacology, and disease. Annu Rev Pharmacol Toxicol, 50, 295–322.

57. Nobrega-Pereira, S., Kessaris, N., Du, T., Kimura, S., Anderson, S. A. & Marin, O. 2008. Postmitotic Nkx2-1 controls the migration of telencephalic interneurons by direct repression of guidance receptors. Neuron, 59, 733–45.

58. O’hare, J. K., Ade, K. K., Sukharnikova, T., Van Hooser, S. D., Palmeri, M. L., Yin, H. H. & Calakos, N. 2016. Pathway-Specific Striatal Substrates for Habitual Behavior. Neuron, 89, 472–9.

59. Okada, K., Nishizawa, K., Fukabori, R., Kai, N., Shiota, A., Ueda, M., Tsutsui, Y., Sakata, S., Matsushita, N. & Kobayashi, K. 2014. Enhanced flexibility of place discrimination learning by targeting striatal cholinergic interneurons. Nat Commun, 5, 3778.

60. Okada, K., Nishizawa, K., Setogawa, S., Hashimoto, K. & Kobayashi, K. 2018. Task-dependent function of striatal cholinergic interneurons in behavioural flexibility. Eur J Neurosci, 47, 1174–1183.

61. Perrin, E. & Venance, L. 2019. Bridging the gap between striatal plasticity and learning. Curr Opin Neurobiol, 54, 104–112.

62. Prado, V. F., Janickova, H., Al-Onaizi, M. A. & Prado, M. A. 2017. Cholinergic circuits in cognitive flexibility. Neuroscience, 345, 130–141.

63. Reiner, A. & Levitz, J. 2018. Glutamatergic Signaling in the Central Nervous System: Ionotropic and Metabotropic Receptors in Concert. Neuron, 98, 1080–1098.

64. Robbe, D. 2018. To move or to sense? Incorporating somatosensory representation into striatal functions. Curr Opin Neurobiol, 52, 123–130.

65. Saunders, A., Granger, A. J. & Sabatini, B. L. 2015. Corelease of acetylcholine and Gaba from cholinergic forebrain neurons. Elife, 4.

66. Straub, C., Saulnier, J. L., Begue, A., Feng, D. D., Huang, K. W. & Sabatini, B. L. 2016. Principles of Synaptic Organization of Gabaergic Interneurons in the Striatum. Neuron, 92, 84–92.

67. Tepper, J. M., Koos, T., Ibanez-Sandoval, O., Tecuapetla, F., Faust, T. W. & Assous, M. 2018. Heterogeneity and Diversity of Striatal Gabaergic Interneurons: Update 2018. Front Neuroanat, 12, 91.

68. Threlfell, S., Lalic, T., Platt, N. J., Jennings, K. A., Deisseroth, K. & Cragg, S. J. 2012. Striatal dopamine release is triggered by synchronized activity in cholinergic interneurons. Neuron, 75, 58–64.

69. Willardsen, M., Hutcheson, D. A., Moore, K. B. & Vetter, M. L. 2014. The Ets transcription factor Etv1 mediates Fgf signaling to initiate proneural gene expression during Xenopus laevis retinal development. Mech Dev, 131, 57–67.

70. Wilson, C. J. & Goldberg, J. A. 2006. Origin of the slow afterhyperpolarization and slow rhythmic bursting in striatal cholinergic interneurons. J Neurophysiol, 95, 196–204.

71. Xu, M., Kobets, A., Du, J. C., Lennington, J., Li, L., Banasr, M., Duman, R. S., Vaccarino, F. M., Dileone, R. J. & Pittenger, C. 2015. Targeted ablation of cholinergic interneurons in the dorsolateral striatum produces behavioral manifestations of Tourette syndrome. Proc Natl Acad Sci U S A, 112, 893–8.

72. Yin, H. H., Knowlton, B. J. & Balleine, B. W. 2004. Lesions of dorsolateral striatum preserve outcome expectancy but disrupt habit formation in instrumental learning. Eur J Neurosci, 19, 181–9.

73. Zhang, Y. F., Reynolds, J. N. J. & Cragg, S. J. 2018. Pauses in Cholinergic Interneuron Activity Are Driven by Excitatory Input and Delayed Rectification, with Dopamine Modulation. Neuron, 98, 918–925 e3.

74. Zhao, Y., Marin, O., Hermesz, E., Powell, A., Flames, N., Palkovits, M., Rubenstein, J. L. & Westphal, H. 2003. The Lim-homeobox gene Lhx8 is required for the development of many cholinergic neurons in the mouse forebrain. Proc Natl Acad Sci U S A, 100, 9005–10.

75. Zhao, Z., Zhang, K., Liu, X., Yan, H., Ma, X., Zhang, S., Zheng, J., Wang, L. & Wei, X. 2016. Involvement of Hcn Channel in Muscarinic Inhibitory Action on Tonic Firing of Dorsolateral Striatal Cholinergic Interneurons. Front Cell Neurosci, 10, 71.

76. Zucca, S., Zucca, A., Nakano, T., Aoki, S. & Wickens, J. 2018. Pauses in cholinergic interneuron firing exert an inhibitory control on striatal output in vivo. Elife, 7.

